# A new technology for isolating organellar membranes provides fingerprints of lipid bilayer stress

**DOI:** 10.1101/2022.09.15.508072

**Authors:** John Reinhard, Leonhard Starke, Christian Klose, Per Haberkant, Henrik Hammarén, Frank Stein, Ofir Klein, Charlotte Berhorst, Heike Stumpf, James P. Sáenz, Jochen Hub, Maya Schuldiner, Robert Ernst

## Abstract

Biological membranes have a stunning ability to adapt their composition in response to physiological stress and metabolic challenges. Little is known how such perturbations affect individual organelles in eukaryotic cells. Pioneering work provided insights into the subcellular distribution of lipids, but the composition of the endoplasmic reticulum (ER) membrane, which also crucially regulates lipid metabolism and the unfolded protein response, remained insufficiently characterized. Here we describe a method for purifying organellar membranes from yeast, MemPrep. We demonstrate the purity of our ER preparations by quantitative proteomics and document the general utility of MemPrep by isolating vacuolar membranes. Quantitative lipidomics establishes the lipid composition of the ER and the vacuolar membrane. Our findings have important implications for understanding the role of lipids in membrane protein insertion, folding, and their sorting along the secretory pathway. Application of the combined preparative and analytical platform to acutely stressed cells reveals dynamic ER membrane remodeling and establishes molecular fingerprints of lipid bilayer stress.

## Introduction

Biological membranes are complex assemblies of lipids and proteins. Their compositions and properties are dynamically regulated in response to stress and various physical and metabolic cues (Harayama & Riezman, 2018; Ernst *et al*, 2018). A prominent example is the homeoviscous adaptation, where the lipid composition is adapted to temperature to maintain membrane fluidity and membrane phase behavior (Sinensky, 1974; Ernst *et al*, 2016; Harayama & Riezman, 2018). Even mammals, which maintain a constant body temperature, can readily adjust their membrane composition in response to dietary perturbation with major impact on collective bilayer properties such as fluidity, thickness, surface charge or stiffness (Bigay & Antonny, 2012; Levental *et al*, 2020). Eukaryotic cells face the challenge of maintaining the properties of not just a single plasma membrane but that of several coexisting organellar membranes each with unique lipid compositions and each exchanging membrane material with other organelles via vesicular carriers and/or lipid transfer proteins. Despite recent advances to manipulate and follow membrane properties (John Peter *et al*, 2022; Renne *et al*, 2022; preprint: Jiménez-Rojo *et al*, 2022; preprint: Tsuchiya *et al*, 2022), we know little about how stressed cells coordinate membrane adaptation between organelles whilst maintaining organelle identity and functions.

The endoplasmic reticulum (ER) is a hotspot for lipid biosynthesis (Zinser *et al*, 1991; Henry *et al*, 2012) and provides an entry to the secretory pathway for soluble and transmembrane proteins. The flux of proteins and lipids through the secretory pathway is controlled by the unfolded protein response (UPR) (Travers *et al*, 2000; Walter & Ron, 2011). When the protein folding capacity of the ER is overwhelmed, unfolded proteins are sensed by the type I membrane protein Ire1 thereby triggering the UPR. The UPR lowers the rate of global protein synthesis, whilst upregulating the ER luminal folding machinery, ER-associated protein degradation, and lipid biosynthesis enzymes (Travers *et al*, 2000; Walter & Ron, 2011). We have recently shown that Ire1 in *Saccharomyces cerevisiae* (from here on ‘yeast’) uses a hydrophobic mismatch-based mechanism to sense aberrant stiffening of the ER membrane thereby rendering it responsive to various lipid metabolic perturbations (Halbleib *et al*, 2017). Known conditions of this membrane-based UPR, termed lipid bilayer stress, include inositol-depletion (Cox *et al*, 1997; Promlek *et al*, 2011), increased lipid saturation (Pineau *et al*, 2009; Surma *et al*, 2013), increased sterol levels (Feng *et al*, 2003; Pineau *et al*, 2009), misregulated sphingolipid metabolism (Han *et al*, 2010), and a disrupted conversion of phosphatidylethanolamine (PE) to phosphatidylcholine (PC) (Ho *et al*, 2020; Ishiwata-Kimata *et al*, 2022). Even prolonged proteotoxic stresses can activate the UPR via a yet poorly defined, membrane-based mechanism (Promlek *et al*, 2011; Väth *et al*, 2021). While lipid bilayer stress is evolutionary conserved among eukaryotes (Volmer *et al*, 2013; Hou *et al*, 2014; Ho *et al*, 2018; Pérez-Martí *et al*, 2022), its molecular manifestations in the ER membrane with respect to the lipid and protein composition remains unexplored.

Regardless of Ire1’s dual sensitivity for proteotoxic and lipid bilayer stress the mechanism of signal transduction is similar for both conditions: Dimerization/Oligomerization of Ire1 juxtaposes its cytosolic kinase/RNase domains thereby triggering the *trans*-autophosphorylation and the activation of the RNase domain (Walter & Ron, 2011; Väth *et al*, 2021). The subsequent cleavage of the *HAC1* mRNA serves as the committed step for generating the Hac1 transcription factor, which translocates to the nucleus and upregulates hundreds of UPR-target genes (Travers *et al*, 2000; Ho *et al*, 2020).

The UPR is a key target of lipotoxicity, and pathological manifestations of chronic ER stress in complex, metabolic diseases such as diabetes, atherosclerosis, and non-alcoholic fatty liver disease feature characteristic lipid fingerprints (Puri *et al*, 2007; Tabassum *et al*, 2019; Lauber *et al*, 2022). Yet, the contribution of lipid bilayer stress to health and disease remains understudied (Hotamisligil, 2010). Yeast is an ideal model organism to decipher fundamental (patho)mechanisms of the lipid metabolic network, the secretory pathway, membrane traffic, and membrane homeostasis (Kurat *et al*, 2006; Henry *et al*, 2012; Ralph-Epps *et al*, 2021). Recent advances in quantitative lipidomics (Ejsing *et al*, 2009) have provided deep insight into the flexibility and adaptation of the cellular lipidome to various metabolic and physical stimuli in both yeast and mammals (Klose *et al*, 2012; Casanovas *et al*, 2015; Levental *et al*, 2020; Surma *et al*, 2021). However, unless these analytical platforms are paired with powerful techniques for isolating organellar membranes from stressed and unstressed cells, they lack the subcellular resolution, which is essential to understand how lipid metabolism is organized between organelles.

Tremendous efforts have been invested in the characterization of organellar membranes (Zinser & Daum, 1995; Schneiter *et al*, 1999; Klemm *et al*, 2009; Surma *et al*, 2011; Reglinski *et al*, 2020), but we still lack comprehensive and quantitative information on the yeast ER. This is probably because the ER forms extensive membrane contact sites (MCSs) with other organelles, which makes its isolation technically challenging (English & Voeltz, 2013; Scorrano *et al*, 2019). Here, we describe a protocol for the isolation of highly enriched organellar membranes, MemPrep. We demonstrate the utility of MemPrep by the successful isolation of both ER and vacuolar membranes from yeast. An in-depth analysis by quantitative lipidomics reveals that the ER membrane is characterized by a particularly high fraction of unsaturated fatty acyl chains in glycerophospholipids (74.1 mol%). Furthermore, our observations suggest the absence of a sterol gradient in the early secretory pathway and a substantial retrograde flux of complex sphingolipids from the Golgi complex to the ER. By analyzing the lipid composition of the stressed ER, we establish molecular fingerprints of lipid bilayer stress and identify a potential role of anionic lipids as negative regulators of the UPR. Our work provides further evidence for an important role of saturated lipids in UPR activation by affecting membrane thickness and stiffness. The MemPrep approach will be crucial to dissect membrane adaptation to metabolic, proteotoxic, and physical stresses on the organelle level in the future.

## Material and Methods

### Generation of MemPrep library

The C-terminus SWAp Tag (SWAT) library from yeast was used to generate a library with a C-terminal tag as previously published (Meurer *et al*, 2018). In short, a SWAT donor strain (yMS2085) was transformed with a donor plasmid (pMS1134) containing the myc-HRV-FLAG cassette and then SWATted as described. The final library genotype is his3Δ1 leu2Δ0 met15Δ0 ura3Δ0, can1Δ::GAL1pr-SceI-NLS::STE2pr-SpHIS5, lyp1Δ::STE3pr-LEU2, XXX::L3-myc-HRV-3xFLAG-ADH1ter-TEFpr-KanMX-TEFter-L4)]. Once generated, to check that proper integration of the cassette into the genome, random proteins were tested by PCR and SDS-PAGE, confirming their in-frame tagging.

### Fluorescence microscopy

3 μl of a yeast cell suspension (OD_600_ = 50), crude cell lysates or a fraction from the isolation procedure were placed on a thin SCD-(1%)-agarose pad and then covered with a coverslip. Images were acquired on an Axio Observer Z1 equipped with a Rolera em-c2 camera (QImaging) and a Colibri 7 (Zeiss) light source for fluorescence excitation. Using either an EC Plan-Neofluar 100x/1.3 or an EC Plan-Apochromat 63x/1.4 objective in combination with a 1.6x tube lens (Zeiss), GFP fluorescence was excited using a 475 nm LED module and a 38 HE filter (Zeiss). Differential interference contrast (DIC) images were acquired using a translight LED light source. Image contrasts were adjusted using Fiji (Schindelin *et al*, 2012).

### Cell cultivation

Cells were cultivated at 30 °C in SCD_complete_ medium (0.79 g/l complete supplement mixture [Formedium, batch no: FM0418/8363, FM0920/10437], 1.9 g/l yeast nitrogen base without amino acids and without ammonium sulfate (YNB) [Formedium, batch no: FM0A616/006763, FM0718/8627], 5 g/l ammonium sulfate [Carl Roth] and 20 g/l glucose [ACS, anhydrous, Carl Roth]) and constantly agitated by shaking the cultures at 220 rpm. Unless stated otherwise, overnight cultures (21 h) were used to inoculate a main culture to an OD_600_ of 0.1. Cells were harvested by centrifugation (3,000 x *g*, 5 min, RT) at an OD_600_ of 1.0, washed with 25 ml ice-cold PBS, snap-frozen in liquid nitrogen, and stored at - 80 °C until further use. In each case ER and vacuolar membranes were isolated from a total of 2,000 and 4,000 OD_600_ units, respectively. This general procedure for cell cultivation and harvesting was also employed for stressed cells, with minor adaptations. For isolating the ER from cells before and after the induction of prolonged proteotoxic stresses, the cells were cultivated in the absence of stress to an OD_600_ of 0.8 and a ‘pre-stress’ sample was harvested. The residual culture was supplemented with either 2 mM dithiothreitol (DTT) or 1.5 μg/ml Tunicamycin (TM) and cultivated for another 4 hours prior to harvesting the cells. For isolating the ER from inositol-depleted cells, a first culture was inoculated to an OD_600_ of 0.003 and cultivated overnight to an OD_600_ of 1.2. Cells from this culture were pelleted (3,000 x *g*, 5 min, RT), washed twice with 100 ml pre-warmed inositol-free medium (SCD_complete-ino_ prepared using yeast nitrogen base lacking inositol (YNB-ino) [Formedium batch no: FM0619/9431]). The washed cells were then resuspended in either inositol-containing SCD_complete_ (control) or in SCD_complete-ino_ (inositol-depletion) medium to an OD_600_ of 0.6 and cultivated for another two hours prior to harvesting the cells.

### Cell lysis and differential centrifugation

Frozen cell pellets of 1,000 OD_600_ units were thawed on ice, resuspended in microsome preparation (MP) buffer (25 mM HEPES pH 7.0, 600 mM mannitol, 1 mM EDTA, 0.03 mg/ml protease inhibitor cocktail [pepstatin, antipain, chymotrypsin] and 12.5 units/ml benzonase nuclease [Sigma Aldrich]) and mechanically disrupted in 15 ml reaction tubes previously loaded with 13 g zirconia/silica beads (0.5 mm diameter, Carl Roth) using a FastPrep-24 bead beater (5 m/s, 10 cycles of 15 s beating and 45 s of cooling in an ice bath). Cell lysates were centrifuged twice (3,234 x *g*, 5 min, 4 °C) in a swinging bucket rotor to remove unbroken cells, cell debris and nuclei. The resulting post nuclear supernatant (PNS) was centrifuged (12,000 x *g*, 20 min, 4 °C) in a Beckman type 70 Ti rotor to remove large organelle fragments. Using the same rotor, the resulting supernatant (S12) was centrifuged (100,000 x *g*, 1 h, 4 °C) to obtain microsomes in the pellet. Microsomes were resuspended in 1 ml MP buffer per 1,000 OD_600_ units original cell mass, snap frozen in liquid nitrogen, and stored at -80 °C until further use. For subsequent proteomics analyses of (pre-)stressed cells (Figure 5), microsomes were additionally resuspended in MP buffer containing 200 mM sodium carbonate (pH 10.6) and incubated rotating overhead at 3 rpm and 4 °C for 1 h to remove soluble and membrane-associated proteins. Carbonate-washed microsomes were neutralized by addition of concentrated HCl, sedimented by ultracentrifugation (100,000 x *g*, 1 h, 4 °C), resuspended in 1 ml MP buffer per 1,000 OD_600_ units original cell mass, snap frozen in liquid nitrogen, and stored at -80 °C until further use.

### Immuno-isolation

The entire isolation procedure was performed on ice or at 4 °C. Microsomes were thawed in 1.5 ml reaction tubes and then dissociated using a sonotrode (MS72) on a Bandelin Sonopuls HD 2070 with 50 % power and 10 pulses of each 0.7 s (duty cycle 0.7). After sonication, the microsomes were centrifuged (3,000 x *g*, 3 min, 4 °C). 700 μl of the resulting supernatant (corresponding to 700 OD_600_ units) were mixed with 700 μl immunoprecipitation (IP) buffer (25 mM HEPES pH 7.0, 150 mM NaCl, 1 mM EDTA) and loaded onto magnetic beads (dynabeads, protein G, 2.8 μm diameter, Invitrogen), which were previously decorated with sub-saturating quantities of a monoclonal anti-FLAG antibody (M2, F1804, Sigma Aldrich). Specifically, the affinity matrix was prepared by using 800 μl of magnetic bead slurry per 700 OD_600_ units of cells, which were incubated overnight at 4 °C with 3.2 μg of the anti-FLAG antibody using an overhead rotor at 20 rpm. Subsequently, microsomes were loaded on the antibody-decorated magnetic beads and allowed to bind for two hours at 4 °C using an overhead rotor at 3 rpm.

The bound membrane vesicles were washed two times with 1.4 ml of wash buffer (25 mM HEPES pH 7.0, 75 mM NaCl, 600 mM urea, 1 mM EDTA) and twice with 1.4 ml of IP buffer. Specific elution was performed by resuspension of the affinity matrix in 700 μl elution buffer (PBS pH 7.4, 0.5 mM EDTA, 1 mM DTT, and 0.04 mg/ml affinity purified GST-HRV3C protease) per 700 OD_600_ units of original cell mass followed by an incubation for two hours at 4 °C on an overhead rotor at 3 rpm. The eluate was centrifuged (264,360 x *g*, 2 h, 4°C) in a Beckman TLA 100.3 rotor to harvest the ER-or vacuole-derived vesicles. The membrane pellet was resuspended in 200 μl PBS per 1,400 OD_600_ units of original cell mass (isolate), snap frozen, and stored at -80 °C until lipid extraction and lipidomics analysis. For proteomics experiments the membrane pellet was resuspended in 40 μl PBS-SDS (1 %) per 1,400 OD_600_ units.

### Liposome fusion assay

POPC liposomes containing 2 mol% 1,2-dioleoyl-sn-glycero-3phosphoethanolamineN-(7-nitro-2-1,3-benzoxadiazol-4-yl) (NBD-PE) and 2 mol% 1,2-dioleoyl-sn-glycero-3-phosphoethanolamine-N-(lissamine rhodamine B sulfonyl) (Rho-PE) were prepared by rehydrating dried lipids in MP buffer. Liposomes were made unilamellar by consecutive extrusion through 0.4 μm and 0.2 μm filters with 13 strokes each. NBD-PE and Rho-PE at a concentration of 2 mol% each form an efficient FRET pair (Weber *et al*, 1998). Labeled liposomes were mixed with an 8.2-fold excess (microsome concentration determined by scattering as described here (Kaiser *et al*, 2011), of unlabelled P100 microsomes and sonicated as described for the immuno-isolation (sonotrode MS72 using a Bandelin Sonopuls HD 2070 with 50 % volume for 10 s as 70 % pulse). Donor fluorescence intensity of the FRET pair in the microsomes-liposomes-mixture (I_DA_) was measured at 530 nm using an Infinite 200 Pro (Tecan) plate reader. Donor only fluorescence (I_D_) was measured at the same wavelength after addition of Triton X-100 to a final concentration of 1 %. Relative FRET efficiencies (E_rel_) were calculated as follows: E_rel_ = 1-(I_DA_/I_D_).

### Lipid extraction for mass spectrometry lipidomics

Mass spectrometry-based lipid analysis was performed by Lipotype GmbH (Dresden, Germany) as described (Ejsing *et al*, 2009; Klose *et al*, 2012). Lipids were extracted using a two-step chloroform/methanol procedure (Ejsing *et al*, 2009). Samples were spiked with internal lipid standard mixture containing: CDP-DAG 17:0/18:1, ceramide 18:1;2/17:0 (Cer), diacylglycerol 17:0/17:0 (DAG), lyso-phosphatidate 17:0 (LPA), lyso-phosphatidylcholine 12:0 (LPC), lyso-phosphatidylethanolamine 17:1 (LPE), lyso-phosphatidylinositol 17:1 (LPI), lyso-phosphatidylserine 17:1 (LPS), phosphatidate 17:0/14:1 (PA), phosphatidylcholine 17:0/14:1 (PC), phosphatidylethanolamine 17:0/14:1 (PE), phosphatidylglycerol 17:0/14:1 (PG), phosphatidylinositol 17:0/14:1 (PI), phosphatidylserine 17:0/14:1 (PS), ergosterol ester 13:0 (EE), triacylglycerol 17:0/17:0/17:0 (TAG), stigmastatrienol, inositolphosphorylceramide 44:0;2 (IPC), mannosyl-inositolphosphorylceramide 44:0;2 (MIPC) and mannosyl-di-(inositolphosphoryl)ceramide 44:0;2 (M(IP)2C). After extraction, the organic phase was transferred to an infusion plate and dried in a speed vacuum concentrator. 1st step dry extract was re-suspended in 7.5 mM ammonium acetate in chloroform/methanol/propanol (1:2:4, V:V:V) and 2nd step dry extract in 33 % ethanol solution of methylamine in chloroform/methanol (0.003:5:1; V:V:V). All liquid handling steps were performed using Hamilton Robotics STARlet robotic platform with the Anti Droplet Control feature for organic solvents pipetting.

### MS data acquisition for lipidomics

Samples were analyzed by direct infusion on a QExactive mass spectrometer (Thermo Scientific) equipped with a TriVersa NanoMate ion source (Advion Biosciences). Samples were analyzed in both positive and negative ion modes with a resolution of Rm/z=200=280000 for MS and Rm/z=200=17500 for MSMS experiments, in a single acquisition. MS/MS was triggered by an inclusion list encompassing corresponding MS mass ranges scanned in 1 Da increments (Surma *et al*, 2015). Both MS and MSMS data were combined to monitor EE, DAG and TAG ions as ammonium adducts; PC as an acetate adduct; and CL, PA, PE, PG, PI and PS as deprotonated anions. MS only was used to monitor LPA, LPE, LPI, LPS, IPC, MIPC, M(IP)2C as deprotonated anions; Cer and LPC as acetate adducts and ergosterol as protonated ion of an acetylated derivative (Liebisch *et al*, 2006).

### Data analysis and post-processing for lipidomics

Data were analyzed with in-house developed lipid identification software based on LipidXplorer (Herzog *et al*, 2011; Herzog *et al*, 2012). Data post-processing and normalization were performed using an in-house developed data management system. Only lipid identifications with a signal-to-noise ratio >5, and a signal intensity 5-fold higher than in corresponding blank samples were considered for further data analysis.

### Sample preparation for proteomics via LC-MS/MS

Lysates were adjusted to 1 % SDS and a final concentration 1 mg/ml. 5 μg of cell lysates and 10 μg of ER membrane were subjected to an in-solution tryptic digest using a modified version of the Single-Pot Solid-Phase-enhanced Sample Preparation (SP3) protocol (Hughes *et al*, 2014; Moggridge *et al*, 2018). In total three biological replicates were prepared including control, wild-type and mutant derived lysates (n=3). Lysates were added to Sera-Mag Beads (Thermo Scientific, #4515-2105-050250, 6515-2105-050250) in 10 μl 15 % formic acid and 30 μl of ethanol. Binding of proteins was achieved by shaking for 15 min at room temperature. SDS was removed by 4 subsequent washes with 200 μl of 70 % ethanol. Proteins were digested overnight at room temperature with 0.4 μg of sequencing grade modified trypsin (Promega, #V5111) in 40 μl Hepes/NaOH, pH 8.4 in the presence of 1.25 mM TCEP and 5 mM chloroacetamide (Sigma-Aldrich, #C0267). Beads were separated, washed with 10 μl of an aqueous solution of 2 % DMSO and the combined eluates were dried down. Peptides of ER membranes were reconstituted in 10 μl of H2O and reacted for 1 h at room temperature with 80 μg of TMT10plex (Thermo Scientific, #90111) (Werner *et al*, 2014) label reagent dissolved in 4 μl of acetonitrile. Peptides of cell lysates were reconstituted in 10 μl of H2O and reacted for 1 h at room temperature with 50 μg of TMT16pro™ label reagent (Thermo Scientific, #A44521) dissolved in 4 μl of acetonitrile. Excess TMT reagent was quenched by the addition of 4 μl of an aqueous 5 % hydroxylamine solution (Sigma, 438227). Peptides were reconstituted in 0.1 % formic acid, mixed to achieve a 1:1 ratio across all TMT-channels and purified by a reverse phase clean-up step (OASIS HLB 96-well μElution Plate, Waters #186001828BA). Peptides were subjected to an off-line fractionation under high pH conditions (Hughes *et al*, 2014). The resulting 12 fractions were then analyzed by LC-MS/MS on an Orbitrap Fusion Lumos mass spectrometer.

### LC-MS/MS analysis of ER membranes

Peptides were separated using an Ultimate 3000 nano RSLC system (Dionex) equipped with a trapping cartridge (Precolumn C18 PepMap100, 5 mm, 300 μm i.d., 5 μm, 100 Å) and an analytical column (Acclaim PepMap 100. 75 × 50 cm C18, 3 mm, 100 Å) connected to a nanospray-Flex ion source. The peptides were loaded onto the trap column at 30 μl per min using solvent A (0.1 % formic acid) and eluted using a gradient from 2 to 38 % Solvent B (0.1 % formic acid in acetonitrile) over 52 min and then to 80 % at 0.3 μl per min (all solvents were of LC-MS grade). The Orbitrap Fusion Lumos was operated in positive ion mode with a spray voltage of 2.4 kV and capillary temperature of 275 °C. Full scan MS spectra with a mass range of 375–1500 m/z were acquired in profile mode using a resolution of 60,000 with a maximum injection time of 50 ms, AGC operated in standard mode and a RF lens setting of 30 %.

Fragmentation was triggered for 3 s cycle time for peptide like features with charge states of 2– 7 on the MS scan (data-dependent acquisition). Precursors were isolated using the quadrupole with a window of 0.7 m/z and fragmented with a normalized collision energy of 36 %. Fragment mass spectra were acquired in profile mode and a resolution of 30,000 in profile mode. Maximum injection time was set to 94 ms or an AGC target of 200 %. The dynamic exclusion was set to 60 s.

### LC-MS/MS analysis of cell lysates

Peptides were separated using an Ultimate 3000 nano RSLC system (Dionex) equipped with a trapping cartridge (Precolumn C18 PepMap100, 5 mm, 300 μm i.d., 5 μm, 100 Å) and an analytical column (Acclaim PepMap 100. 75 × 50 cm C18, 3 mm, 100 Å) connected to a nanospray-Flex ion source. The peptides were loaded onto the trap column at 30 μl per min using solvent A (0.1 % formic acid) and eluted using a gradient from 2 to 80 % Solvent B (0.1 % formic acid in acetonitrile) over 2 h at 0.3 μl per min (all solvents were of LC-MS grade). The Orbitrap Fusion Lumos was operated in positive ion mode with a spray voltage of 2.4 kV and capillary temperature of 275 °C. Full scan MS spectra with a mass range of 375–1500 m/z were acquired in profile mode using a resolution of 120,000 with a maximum injection time of 50 ms, AGC operated in standard mode and a RF lens setting of 30 %.

Fragmentation was triggered for 3 s cycle time for peptide like features with charge states of 2– 7 on the MS scan (data-dependent acquisition). Precursors were isolated using the quadrupole with a window of 0.7 m/z and fragmented with a normalized collision energy of 34 %. Fragment mass spectra were acquired in profile mode and a resolution of 30,000 in profile mode. Maximum injection time was set to 94 ms or an AGC target of 200 %. The dynamic exclusion was set to 60 s.

### Data analysis

Acquired data were analyzed using IsobarQuant (Franken *et al*, 2015) and Mascot V2.4 (Matrix Science) using a reverse UniProt FASTA *Saccharomyces cerrevisiae* database (UP000002311) including common contaminants and the following Rtn1-myc-3C-3xFLAG-tagged (bait) protein employed for the enrichment of subcellular membranes:

sp|P1707_RE|P1707_RE MSASAQHSQAQQQQQQKSCNCDLLLWRNPVQTGKYFGGSLLALLILKKVNLITFFLKVAYTILFTTGSIEFVSKLFLGQGLITKYGPKECPNIAGFIKPHIDEALKQLPVFQAHIRKTVFAQVPKHTFKTAVALFLLHKFFSWFSIWTIVFVADIFTFTLPVIYHSYKHEIDATVAQGVEISKQKTQEFSQMACEKTKPYLDKVESKLGPISNLVKSKTAPVSSTAGPQTASTSKLAADVPLEPESKAYTSSAQVMPEVPQHEPSTTQEFNVDELSNE LKKSTKNLQNELEKNNAGGGGGGEQKLISEEDLGSGLEVLFQGPGSGDYKDHDGDYKDHDIDYKDD DDK

The following modifications were taken into account: Carbamidomethyl (C, fixed), TMT10plex (K, fixed), Acetyl (N-term, variable), Oxidation (M, variable) and TMT10plex (N-term, variable). TMT16plex labeled samples The TMT16plex (K, fixed) and TMT16plex (N-term, variable) labels were considered as modifications. The mass error tolerance for full scan MS spectra was set to 10 ppm and for MS/MS spectra to 0.02 Da. A maximum of 2 missed cleavages were allowed. A minimum of 2 unique peptides with a peptide length of at least seven amino acids and a false discovery rate below 0.01 were required on the peptide and protein level (Savitski *et al*, 2015).

### ER enrichment calculation based on untargeted proteomics

IsobarQuant output data were analyzed on a gene symbol level in R (https://www.R-project.org) using in-house data analysis pipelines. In brief, data was filtered to remove contaminants and proteins with less than 2 unique quantified peptide matches. Subsequently, protein reporter signal sums were normalized within the TMT set using the vsn package (Huber *et al*, 2002) and fold changes were calculated over vsn-corrected values in the total lysate channel of the respective replicate. Gene ontology (GO) term annotations were retrieved from Uniprot (accessed 8.3.2021) (Supplementary Table S1). Compartment-specific unique annotations were obtained by aggregating the following GO terms: GO:0005576, GO:0031012 to *extracellular region*; GO:0005886 to *plasma membrane*; GO:0005737, GO:0005829 to *cytoplasm*; GO:0005739 to *mitochondrion*; GO:0005777 to *peroxisome*; GO:0005783 to *ER*; GO:0005794 to *Golgi apparatus*; GO:0005634, GO:0005654, GO:0005730 to *nucleus*; GO:0031965 to *nuclear membrane*; GO:0005773, GO:0000324, GO:0005774 to *vacuole*; and GO:0005811 to *lipid droplet*.

### Molecular dynamics (MD) simulations

All-atom MD simulations were set up and carried out using the GROMACS software (Páll *et al*, 2020). Lipid topologies and structures were taken from the CHARMM-GUI web server (Jo *et al*, 2009). Bilayers were then generated using MemGen (Schott-Verdugo & Gohlke, 2019). Three different ER compositions were used (Supplementary Table S2) as well as a reference membrane composed of 50 % POPC and 50 % DOPC. Each system contained 100 lipids per leaflet and 60 water molecules per lipid. Na^+^ and Cl^-^ ions were added to reach an ionic concentration of 0.15 M. Taken together each system contained approximately 60000 atoms. Simulations were carried out using the CHARMM36m forcefield (Huang *et al*, 2017) and the CHARMM-modified TIP3P water model (Jorgensen *et al*, 1998). The system was kept at 303 K using velocity-rescaling (Bussi *et al*, 2007). Semi-isotropic pressure coupling at 1.0 bar was applied using the Berendsen barostat (Berendsen *et al*, 1984) for equilibration and the Parrinello-Rahman barostat using a coupling time constant of τ = 2 ps (Parrinello & Rahman, 1981) during production runs. Electrostatic interactions were calculated using the particle-mesh Ewald method (Essmann *et al*, 1995). A cutoff of 1.2 nm according to CHARMM36 specifications (Best *et al*, 2012) was used for the non-bonded interactions, while the Lennard-Jones interactions were gradually switched off between 1.0 and 1.2 nm. Bond constraints involving hydrogen atoms were implemented using LINCS (Hess, 2008), thus a 2 fs time step was chosen. Each system was initially equilibrated for 50 ns followed by three independent production runs of at least 700 ns.

For the analysis, the first 100 ns of each production run was discarded. Mass density profiles were calculated along the z-axis (membrane normal) using the Gromacs module gmx density. The membrane thickness was extracted from the density profiles using a threshold of 500 kg/m^3^. Errors were obtained by averaging and calculating the SEM over independent simulations segments. An in-house modified version of the Gromacs module freevolume was used to calculate the free volume profile as a function of z. The module estimates the accessible free volume by inserting probe spheres of radius R at random positions, while testing the overlap with the Van der Waals radii of all simulated atoms. Here, we used a probe radius or R=0, thereby obtaining the total free volume. Errors for each individual simulation were obtained by multiple independent insertion rounds carried out by gmx freevolume, the overall error for the averaged curves was calculated using standard error propagation. Surface packing defects were calculated using the program PackMem (Gautier *et al*, 2018). The program uses a grid-based approach to identify surface defects and distinguishes between deep and shallow defects based on the distance to the mean glycerol position (everything below a threshold of 1 Å was considered a deep defect). By fitting a single exponential to the obtained defect area distribution, the defect size constant was determined for each defect type. To achieve converged results, trajectory snapshots taken every 100 ps were used. Error estimation was conducted by block averaging dividing each simulation into 3 blocks of equal size and calculating the SEM over all blocks.

### RNA preparation, cDNA synthesis, and RT-qPCR analysis

UPR activation was measured by determining the mRNA levels of spliced *HAC1, PDI*, and *KAR2*. For each experimental condition total RNA was prepared from 5 OD_600_ units of cells using the RNeasy Plus RNA Isolation Kit (Qiagen). The synthesis of cDNA was performed using 500 ng of prepared RNA, Oligo(dT) primers, and the Superscript II RT protocol (Invitrogen).

RT-qPCR was performed using the ORA qPCR Green ROX L Mix (HighQu) and a Mic qPCR cycler (Bio Molecular Systems) in a 20 μl reaction volume. Following primers were used at a final concentration of 400 nM to determine the CT values of the housekeeping gene *TAF10* and genes of interest: spliced *HAC1* forward: 5’-TACCTGCCGTAGACAACAAC-3’; spliced *HAC1* reverse: 5’-ACTGCGCTTCTGGATTAC-3’; *PDI* forward: 5’-TTCCCTCTATTTGCCATCCAC-3’; *PDI* reverse: 5’-GCCTTAGACTCCAACACGATC-3’; *KAR2* forward: 5’-TGCTGTCGTTACTGTTCCTG-3’; *KAR2* reverse: 5’-GATTTATCCAAACCGTAGGCAATG-3’; *TAF10* forward: 5’-ATATTCCAGGATCAGGTCTTCCGTAGC-3’; *TAF10* reverse: 5’-GTAGTCTTCTCATTCTGTTGATGTTGTTGTTG-3’.

The qPCR program consisted of the following steps: (1) 95 °C, 15 min; (2) 95 °C, 20 s; (3) 62 °C, 20 s; (4) 72 °C, 30 s; and (5) 72 °C, 5 min; steps 2–4 were repeated 40 times. We used the comparative ΔΔCT method with normalization to *TAF10* CT values to quantify the levels of spliced *HAC1, PDI*, and *KAR2* mRNA.

## Results

### Creation of a rapid and clean approach for yeast organelle isolation, MemPrep

In the past organelle isolation in yeast has been carried out predominantly by differential sedimentation and density centrifugation (Zinser & Daum, 1995; Schneiter *et al*, 1999). Affinity purification methods that work well in mammalian cells cannot be translated easily into yeast work, especially when the organelle-of-interest forms extensive membrane contact sites (Takamori *et al*, 2006; Klemm *et al*, 2009; Surma *et al*, 2011; Abu-Remaileh *et al*, 2017). We sought to create a yeast-specific affinity purification method for obtaining clean organelle fractions, MemPrep. We reasoned that important aspects of MemPrep would be the capacity to rapidly bind organellar membranes with high specificity and the ability to release them selectively after intense washing. Hence, we constructed a tagging-cassette that can equip an open reading frame in yeast with a sequence encoding for a C-terminal bait tag comprising a myc epitope, a recognition site for the human rhinovirus (HRV) 3C protease, and three repeats of the FLAG epitope (Figure 1A). Following proof of concept of the validity of MemPrep (see below) and to enable our approach to be widely used by the yeast community regardless of which organelle is of interest, we created a systematic collection of strains in which every yeast protein is tagged with the bait sequence (see some examples for each organelle in Supplementary Figure S1A). To do this we used the SWAp Tag (SWAT) approach (Yofe *et al*, 2016; Meurer *et al*, 2018; Weill *et al*, 2018) coupled with automated library creation strategies (Tong & Boone, 2006; Weill *et al*, 2018). The library or any individual strain are freely distributed.

**Figure 1.**
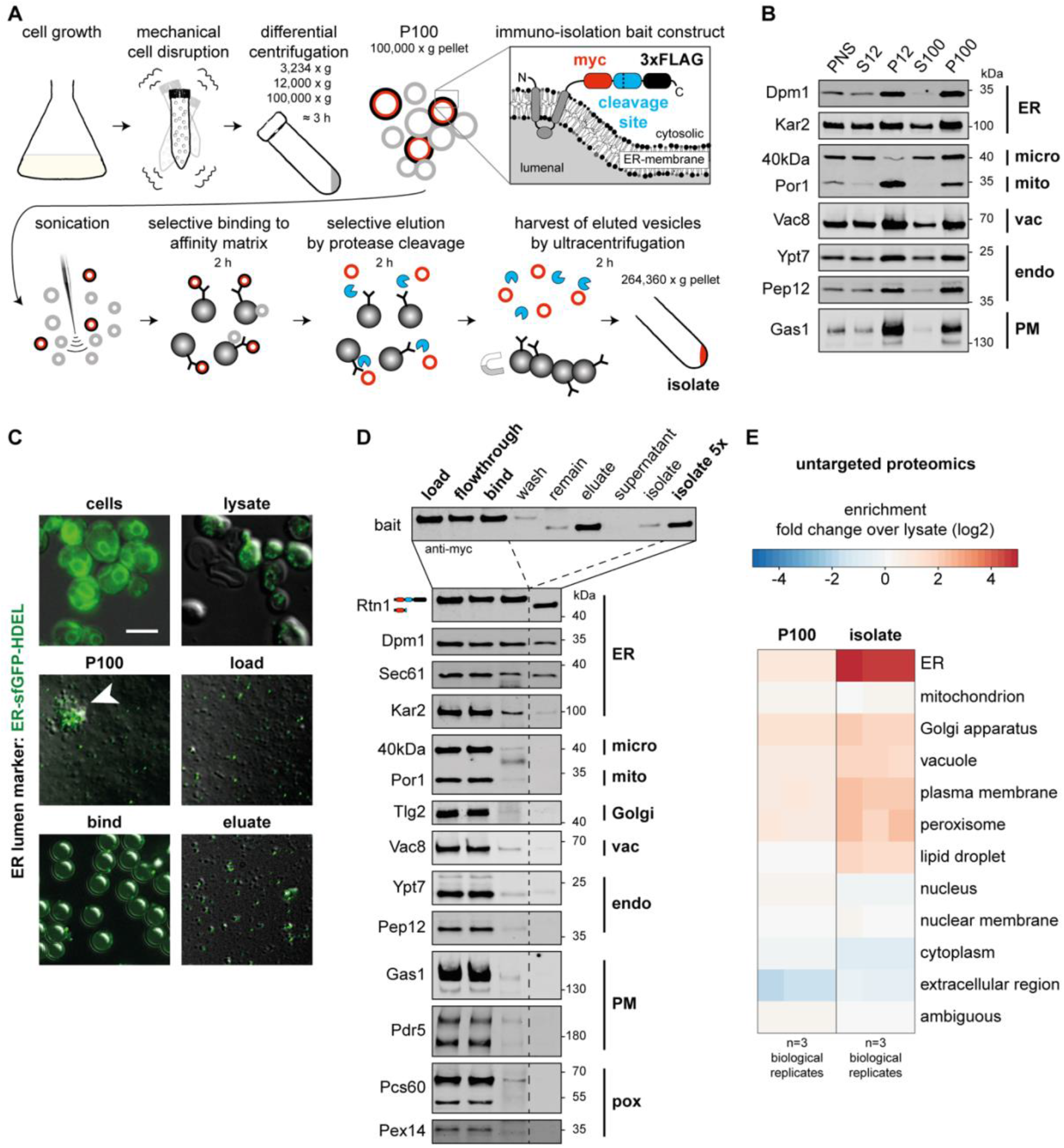
Immuno-isolation of the ER via MemPrep. **(A)** Schematic representation of the immuno-isolation protocol. Cells are cultivated in SCD_complete_ medium and mechanically disrupted by vigorous shaking with zirconia/glass beads. Differential centrifugation at 3,234 x g, 12,000 x g, and 100,000 x g yields crude microsomes in the P100 fraction originating from different organelles. The bait tag installed at the C-terminal end of Rtn1 for the immuno-isolation is depicted in the inlay (myc-tag, human rhinovirus (HRV) 3C protease cleavage site, 3xFLAG-tag). Sonication segregates clustered vesicles and lowers the vesicle size. ER-derived vesicles are specifically captured by anti-FLAG antibodies bound to Protein-G on magnetic beads. After rigorous washing, the ER-derived vesicles are selectively eluted by cleaving the bait tag with the HRV 3C protease (blue sectors). The eluted ER-derived vesicles (red circles) are harvested and concentrated by ultracentrifugation. **(B)** Distribution of the indicated organellar markers in the fractions of a differential centrifugation procedure: Supernatant after 3,234 x g centrifugation (post-nuclear supernatant, PNS), supernatant after 12,000 x g centrifugation (S12), pellet after 12,000 x g centrifugation (P12), supernatant after 100,000 x g centrifugation (S100), pellet after 100,000 x g centrifugation (P100). Dpm1 and Kar2 are ER markers, the 40 kDa protein (40kDa) is a marker for light microsomes (Zinser *et al*, 1991), Por1 is a marker of the outer mitochondrial membrane, Vac8 is a vacuolar marker, Ypt7 and Pep12 mark endosomes, and Gas1 serves as plasma membrane marker. 7.8 μg total protein loaded per lane. **(C)** Overlay of fluorescence micrographs and differential interference contrast images of cells and isolation fractions containing an ER luminal marker (ER-sfGFP-HDEL). Intact cells (cells) show typical ER staining. Mechanical cell disruption leads to fragmentation and release of intracellular membranous organelles (lysate). The crude microsomal fraction (P100) contains aggregates of GFP-positive and GFP-negative vesicles (white arrowhead). Segregation by sonication yields more homogenous size distribution of vesicles (load). Individual ER luminal marker containing vesicles are bound to the surface of much larger magnetic beads (bind). Selective elution by protease cleavage releases vesicles from the affinity matrix (eluate). **(D)** Immunoblot analysis of immuno-isolation fractions for common organellar markers (ER, endoplasmic reticulum; micro, microsomal fraction; mito, mitochondria; Golgi, Golgi apparatus; vac, vacuole; endo, endosomal system; PM, plasma membrane; pox, peroxisomes). 0.2 % of each fraction loaded per lane. **(E)** Untargeted protein mass spectrometry analysis showing enrichment of P100 and isolate fractions over whole cell lysate. The determination of organelle enrichment of proteins is based on uniquely annotated cellular compartment gene ontology terms.

### MemPrep yields highly enriched ER membrane vesicles

To showcase MemPrep we first focused on the largest organelle in the cell, the ER, which is a particularly challenging target. It forms physical contact sites with almost every other membrane-bound organelle (English & Voeltz, 2013) and previous attempts to isolate ER membranes suffered from significant mitochondrial contaminations (Schneiter *et al*, 1999; Reglinski *et al*, 2020). We used Rtn1 as a bait for ER proteins since it is a small and highly abundant reticulon protein (∼37,100 copies per cell), which stabilizes membrane curvature in the tubular ER (Ghaemmaghami *et al*, 2003; Voeltz *et al*, 2006). Several experimental factors are important to ensure the successful isolation of ER membranes (Figure 1A): Firstly, cells are mechanically disrupted, thereby minimizing potential artifacts from the ongoing lipid metabolism and ER stress that occurs during the digestion of the cell wall under reducing conditions (Zinser & Daum, 1995; Klemm *et al*, 2009; Reinhard *et al*, 2020). Secondly, large organellar fragments are disrupted by sonication prior to the immuno-isolation because small organellar fragments are less likely to form physical contacts to other organelles. Thirdly, physical membrane contacts between vesicles are destabilized by urea-containing wash buffers. And fourthly, the isolated membrane vesicles are selective eluted from the affinity matrix thereby providing a straightforward coupling to various mass spectrometry-based analytical platforms following previous paradigms (Klemm *et al*, 2009).

Enrichments of organellar membranes relies first on differential centrifugation and only then on an affinity isolation. To decide on the exact fraction best to utilize for membrane pull downs, we performed immunoblotting experiments after differential centrifugation (Figure 1B). Membrane markers for the ER (Dpm1), mitochondria (Por1), endosomes (Ypt7, Pep12), the vacuole (Vac8), and the plasma membrane (Gas1) were all enriched in the pellets of a centrifugation at 12,000 x g (P12) and 100,000 x g (P100), while the light microsomal 40 kDa protein (40kDa) was found predominantly found in the P100 fraction (Figure 1B). The marker for the outer mitochondrial membrane (Por1) was significantly enriched in P12 relative to P100 (Figure 1B). To minimize contaminations from mitochondrial membranes we decided to use the crude microsomal P100 fraction for isolating ER membrane vesicles. Of note, a substantial fraction of the ER-luminal chaperone Kar2 was found in the supernatant after centrifugation at 100,000 x g (S100), hence suggesting that at least some of the ER-luminal contents are released during cell disruption.

To ensure that our choice of P100 is optimal and to uncover the extent of loss of ER luminal proteins we followed the entire process from cell disruption to the elution of the isolated vesicles in a control experiment using fluorescence microscopy. To this end, we used cells expressing not only an ER bait protein, but also an ER-targeted, superfolder-GFP variant equipped with a HDEL sequence for ER retrieval (Figure 1C) (Lajoie *et al*, 2012). By following the fluorescent ER-luminal marker, we realized that the crude microsomal P100 fraction contains clumps of both GFP-positive and GFP-negative vesicles (Figure 1C; P100, white arrowhead). Due to the loss of ER luminal proteins, the GFP-negative vesicles could be derived from the ER but may also be from other organelles. Regardless, clumps of vesicles would make isolation impossible, and hence we decided to separate them using ultrasound (Figure 1A). Indeed, this eliminated the presence of aggregated vesicles (Figure 1C).

Sonication transiently disrupts membranes and might induce fusion of ER membrane vesicles with non-ER membranes. Because this would obscure our measurement of the ER membrane composition, we performed control experiments to rule out this possibility. We utilized small unilamellar vesicles with a fusogenic lipid composition (POPC/DOPE/POPS/Erg/NBD-PE/Rho-PE at 61/20/5/10/2/2 mol%, respectively) that includes two fluorescent lipids forming a Förster resonance energy transfer (FRET) pair. We sonicated these synthetic liposomes in the presence of excess microsomal membranes (P100). Because fusion between the synthetic liposomes and microsomal membranes would ‘dilute’ the fluorescent lipid analogs, a decrease of the relative FRET efficiency would be expected upon mixing of membrane contents. However, the 10 cycles of sonication used in MemPrep for dissociating vesicle aggregates do not cause a substantial membrane mixing. Only after 100 cycles, which also leads to sample warming, or in the presence of Ca^2+^ and PEG, which triggers membrane fusion, we observed lower FRET efficiencies indicative for lipid exchange and membrane fusion (Supplementary Figure S1B). While we expect that some ER-luminal proteins are released during the sonication step, our data exclude the possibility the dissociation of aggregated vesicles causes a significant degree of membrane mixing from fusion and/or lipid exchange.

After having optimized sample homogenization, we turned our attention to the immuno-isolation procedure. We decided on Protein G-coated, magnetic dynabeads decorated with anti-FLAG antibodies at sub-saturating concentrations. A low density of antibodies is required to lower steric hindrances and unwanted avidity effects, which might eventually impede the elution of membrane vesicles from the matrix. The capturing of GFP-positive, ER-derived vesicles to the affinity matrix was validated by fluorescence microscopy (Figure 1C, bind). After extensive washing with urea-containing buffers, the isolated vesicles were eluted (Figure 1C; eluate) by cleaving the bait tag as validated by immunoblotting using anti-myc antibodies (Figure 1D; eluate). The isolated membrane material was harvested and concentrated by ultracentrifugation (264,360 x g, 2 h, 4 °C) (Figure 1A). Immunobloting demonstrated the co-purification of the bait (Rtn1) with other ER membrane proteins (Dpm1, Sec61) (Figure 1D), while most of the ER-luminal chaperone Kar2 is lost during the isolation. Remarkably, all markers for other organelles including the light microsomal marker 40 kDa protein (40kDa) and the mitochondrial marker Por1 were absent in the final isolate (Figure 1D). This shows that the isolation of ER-derived membranes via MemPrep is free of considerable contaminations.

While loss of several specific organelle markers by immunoblotting is often used as a ‘gold standard’, we decided to go one step further. Consequently, we measured the level of cleanliness of our preparations by TMT-multiplexed, untargeted protein mass spectrometry to estimate the enrichment of the ER membrane proteins in the immuno-isolate relative to the cell lysate (Figure 1E). We compared the enrichment of the ER with that of other organelles using a total of 1591 proteins uniquely annotated for cellular compartments with gene ontology terms (GO terms) (Figure 1E, Supplementary Figure S1C, Supplementary Table S1). The median enrichment of 178 ER-specific proteins was 25.7-fold in the immuno-isolate over the cell lysate, which is also consistent with semi-quantitative, immunoblotting data (Supplementary Figure S1D). Most notably, quantitative proteomics approach revealed the efficient depletion of cytosolic (cytoplasm), inner nuclear membrane, and even mitochondrial proteins, which represented a major contaminant in microsome preparations in the past (Schneiter *et al*, 1999; Reglinski *et al*, 2020). Hence, we have isolated ER-derived membrane vesicles from yeast with unprecedented purity.

### The lipid composition of the ER

Previous attempts of establishing subcellular membrane compositions in yeast have either not included the ER or yielded insufficiently pure preparations (Schneiter *et al*, 1999; Reglinski *et al*, 2020). Having established the isolation of ER-derived membranes, we were interested in determining their lipid composition using state-of-the-art, quantitative lipidomics (Figure 2A-C, Supplementary Table S3). We found that the ER membrane features (compared to the lipid composition of the corresponding cell lysate) 1) substantially lower levels of neutral storage lipids (ergosterol esters (EEs) and triacylglycerols (TAGs)), 2) significantly more diacylglycerol (DAG), phosphatidylcholine (PC) and phosphatidylethanolamine (PE), but 3) less phosphatidylinositol (PI) lipids. Hence, the ER maintains a characteristic lipid composition even though it readily exchanges membrane material with other organelles (Wong *et al*, 2019). Remarkably, the level of ergosterol in the ER (9.7 mol%) is barely distinct from the level in whole cells (10.5 mol%) (Figure 2A) or in the *trans*-Golgi network/endosome (TGN/E) system (9.8 mol%) (Klemm *et al*, 2009), but much lower than in the plasma membrane (>44 mol%) (Surma *et al*, 2011). The absence of a robust sterol gradient in the early secretory pathway has important implications for the sorting of transmembrane proteins (Ridsdale *et al*, 2006; Lorent *et al*, 2017) and is consistent with a direct ER-to-Golgi delivery via membrane bridges (Weigel *et al*, 2021). Furthermore, we find complex sphingolipids such as inositol-phosphorylceramide (IPC), mannosyl-IPC (MIPC), and mannosyl-di-(IP)C (M(IP)_2_C) at considerable levels in the ER (Figure 2B). Because these lipids are synthesized only in the Golgi apparatus, our finding suggests a substantial retrograde flux of complex sphingolipids from the Golgi complex to the ER. It is likely that these lipids are delivered to the ER via COP-I vesicles together with ER-resident proteins bound to the HDEL-receptor (Semenza *et al*, 1990; Aguilera-Romero *et al*, 2008). A closer look at the fatty acyl chain composition of ER lipids reveals a particularly low level (<5 mol%) of tightly-packing, saturated lipids and a significant enrichment of loosely packing, unsaturated lipids (Figure 2C). Loose lipid packing and low membrane rigidity are likely contributing to the remarkable ability of the ER to accept and fold the entire diversity of transmembrane proteins differing substantially in shape and hydrophobic thicknesses (Sharpe *et al*, 2010; Quiroga *et al*, 2013; Lorent *et al*, 2020). Future work will be dedicated to quantifying also phosphorylated PI species such as phosphatidylinositol-4, 5-bisphosphate (PIP2) or phosphatidylinositol-3, 4, 5-triphosphate (PIP3).

**Figure 2.**
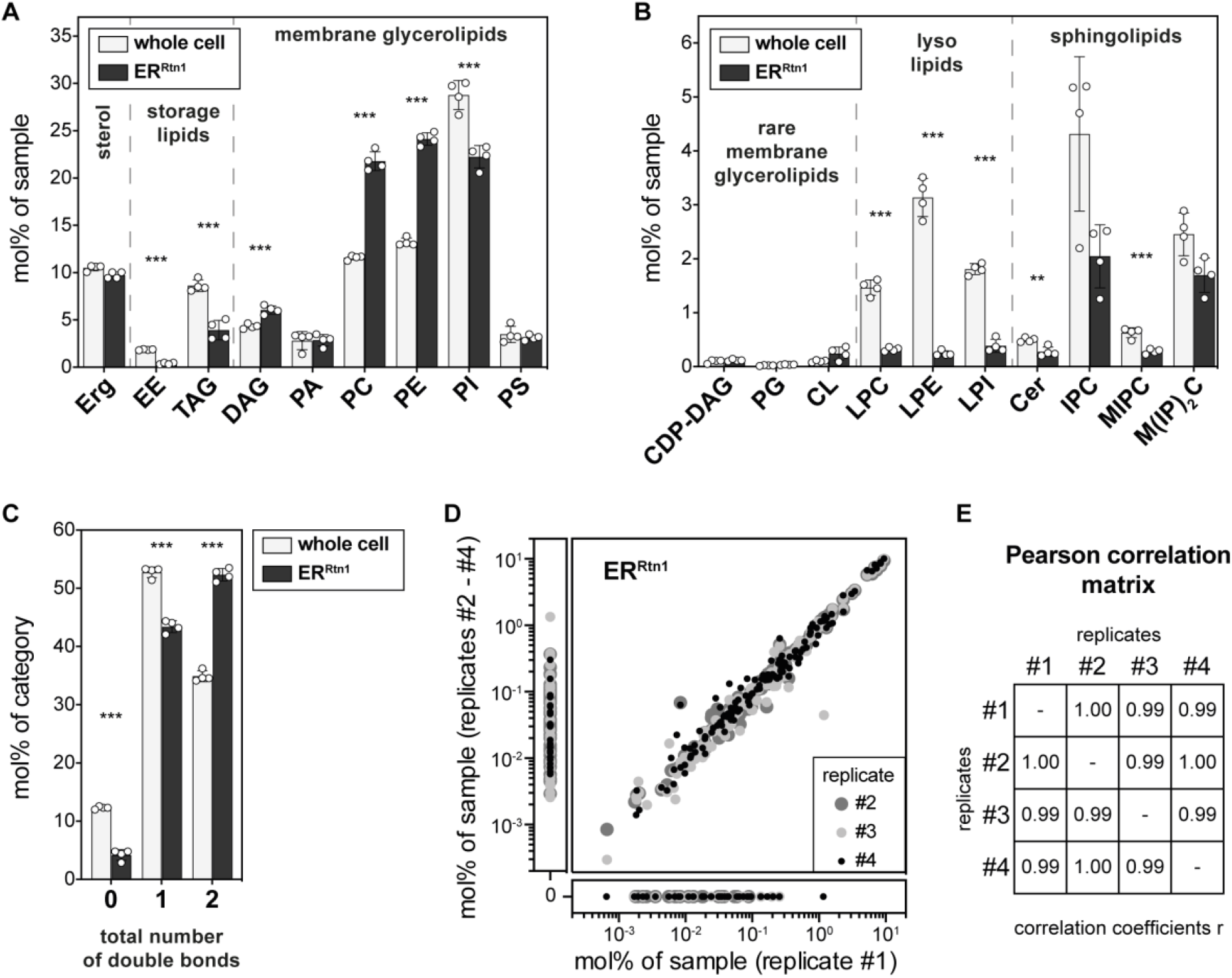
Lipid composition of the ER of *S. cerevisiae*. SCD_complete_ medium was inoculated with Rtn1-bait cells to an OD_600_ of 0.1 from an overnight pre-culture and cells were harvested at an OD_600_ of 1.0. ER derived membranes were purified by differential centrifugation and immuno-isolation and subsequently analyzed by quantitative shotgun lipidomics. **(A)** Lipid class composition given as mol% of all lipids in the sample. Classes are categorized into sterol (Erg, ergosterol), storage lipids (EE, ergosteryl ester; TAG, triacylglycerol), membrane glycerolipids (DAG, diacylglycerol; PA, phosphatidic acid; PC, phosphatidylcholine; PE, phosphatidylethanolamine; PI, phosphatidylinositol; PS, phosphatidylserine). **(B)** Continuation of lipid class composition given as mol% of all lipids in the sample. Classes are categorized into rare membrane glycerolipids (CDP-DAG, cytidine diphosphate diacylglycerol; PG, phosphatidylglycerol; CL, cardiolipin), lysolipids (LPC, lyso-phosphatidylcholine; LPE, lyso-phosphatidylethanolamine; LPI, lyso-phosphatidylinositol) and sphingolipids (Cer, ceramide; IPC, inositolphosphorylceramide; MIPC, mannosyl-IPC; M(IP)_2_C, mannosyl-di-IPC). **(C)** Total number of double bonds in membrane glycerolipids, except CL, (*i*.*e*. CDP-DAG, DAG, PA, PC, PE, PG, PI, PS) as mol% of this category. **(D)** Reproducibility of immuno-isolated ER lipidome data shown as correlation of mol% of sample values of all detected lipid species between replicates 1 and replicates 2-4. **(E)** Pearson correlation coefficients of lipidomics data for all combinations of replicate samples. Statistical significance was tested by multiple t tests correcting for multiple comparisons using the method of Benjamini, Krieger and Yekutieli, with Q = 1 %, without assuming consistent standard deviations. *p < 0.05, **p < 0.01, ***p < 0.001.

In summary, our molecular analysis of the ER membrane reveals several surprising insights, which are nevertheless consistent with our current understanding of the properties and functions of the ER. The robustness and reproducibility of our MemPrep approach coupled to lipidomic platforms is demonstrated by the near perfect correlation of lipid abundances reported in four independent experiments (Figure 2D, E).

### Stable lipid compositions after cell lysis contrasts ER lipid remodeling in living cells

While our isolation process is shorter than many previously employed methods for organelle purification, it still takes 8 h from cell lysis to finish. Hence, we wanted to exclude that ongoing lipid metabolism during the isolation procedure distorts the measured lipid composition. Consequently, we performed a control experiment in which we split a crude microsome preparation (P100) into two equal samples. The first sample was directly snap-frozen in liquid N_2_ while the second one frozen only after an incubation at 4 °C for 8 h. A comparison of the two samples revealed remarkably similar lipid compositions (Supplementary Figure S2A-C, Supplementay Table S3). Only the low abundant lyso-PC, lyso-PE, and lyso-PI lipids showed some differences (Supplementary Figure S2B) thereby suggesting a loss of lysolipids over time consistent with their role as intermediates of lipid degradation (Harayama & Riezman, 2018). The overall stability of the lipidome, however, supports the view that the lipid composition of the immuno-isolated ER membranes reflects the original, *in vivo* lipid composition of the ER.

To investigate the responsiveness of the ER to metabolic perturbation (Zinser *et al*, 1991; Henry *et al*, 2012), we immuno-isolated ER-derived membranes from cells cultured in synthetic complex dextrose (SCD) with or without additional 2 mM choline and determined the resulting lipid composition (Supplementary Figure S3A-C, Supplementary Table S3). Choline is a lipid metabolite that can be activated to CDP-choline and then transferred onto diacylglycerol (DAG) to yield PC (Supplementary Figure S3D) (Kennedy & Weiss, 1956; Henry *et al*, 2012). Somewhat expectedly, the ER of choline-challenged cells features substantially higher levels of PC and lower levels of PE, which is also reflected by an increase of the PC-to-PE ratio from ∼1.1 to ∼2.4 (Supplementary Figure S3A). While the abundance of most other lipid classes including ergosterol, DAG, phosphatidic acid (PA) and IPC are unaffected by the metabolic challenge, we also observe mildly increased abundances of PS and MIPC, and mildly decreased levels of PI. Intriguingly, the increased PC-to-PE ratio of ∼2.4 in choline-challenged cells is not associated with changes in lipid saturation (Supplementary Figure S3C). Plotting the choline-induced changes of the lipid fatty acyl chains reveals only minimal changes: the average chain length of PC is decreasing, while it is increasing for PE, PS, and PI (Supplementary Figure S3E). Even though the metabolic challenge substantially perturbs the PC-to-PE ratio, it does not trigger the UPR, as judged by RT-qPCR experiments quantifying the abundance of the UPR-specific, spliced *HAC1* mRNA and that of the two UPR target genes *KAR2* and *PDI1*. This is in stark contrast to inositol-depletion, which robustly activates the UPR (Supplementary Figure S3F) (Cox *et al*, 1997; Promlek *et al*, 2011). Together, these data highlight the accuracy by which MemPrep can measure even slight differences in organelle lipid composition on the one hand and the remarkable versatility of the lipid modifying enzymes to maintain similar membrane properties despite changes in head-group composition, on the other.

### A molecular fingerprint of lipid bilayer stress during inositol-depletion

Having shown the ability of the ER to adjust its membrane composition, we turned our attention to the stressed ER. Lipid bilayer stress is a collective term for aberrant ER membrane compositions activating the UPR (Ho *et al*, 2018; Radanović & Ernst, 2021). The acute depletion of inositol causes a robust, but transient UPR without triggering a substantial accumulation of misfolded proteins (Supplementary Figure S3F) (Cox *et al*, 1997; Promlek *et al*, 2011; Lajoie *et al*, 2012). By combining MemPrep with a quantitative lipidomics platform, we set out to define, for the first time, the molecular fingerprints of lipid bilayer stress in the ER (Figure 3A, B).

**Figure 3.**
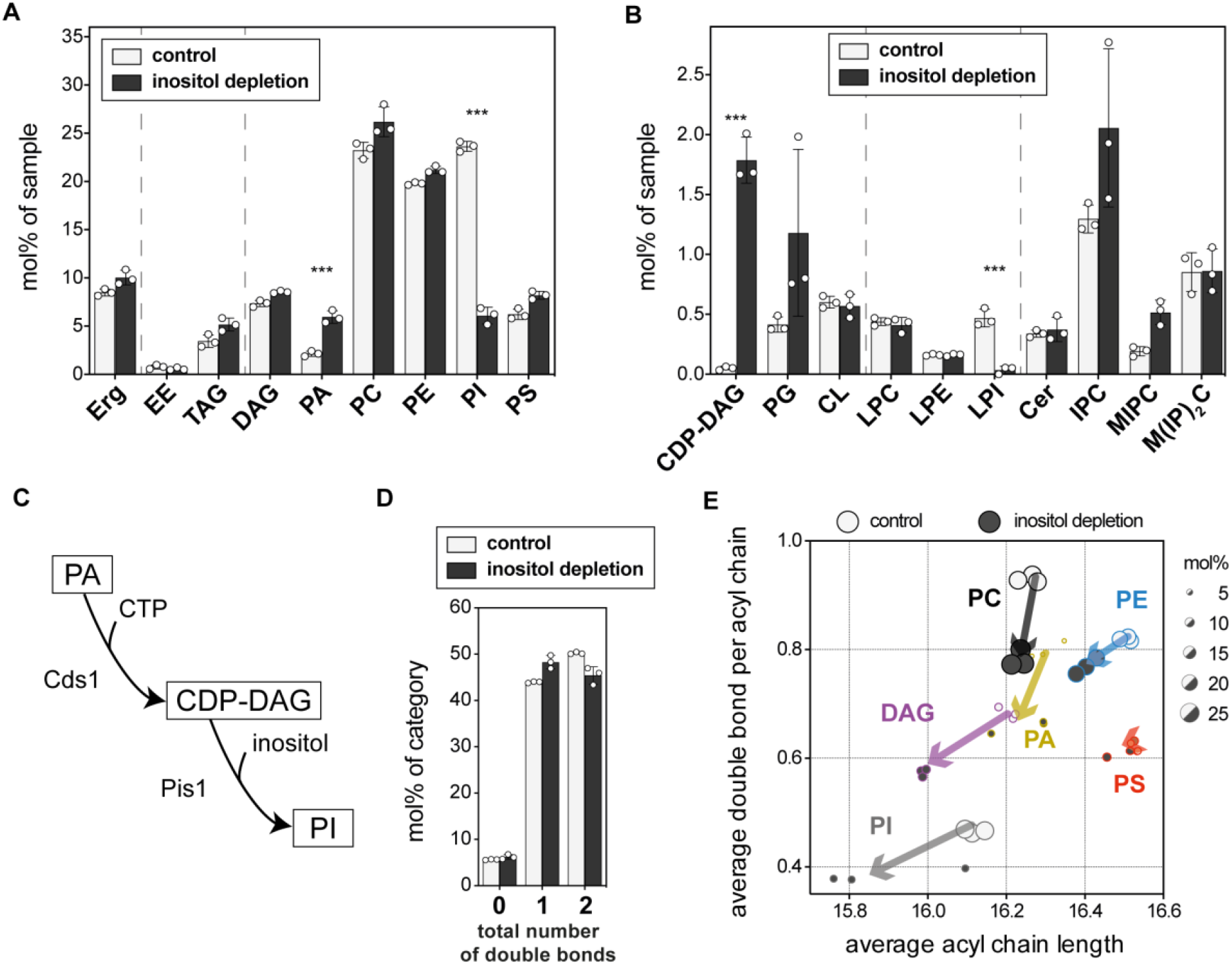
Lipid fingerprints of lipid bilayer stress. SCD_complete_ medium was inoculated with Rtn1-bait cells to an OD_600_ of 0.003 from an overnight pre-culture and grown to an OD_600_ of 1.2. Cells were washed with inositol free medium and then cultivated for an additional 2 h in either inositol-free (inositol depletion) or SCD_complete_ medium (control) starting with an OD_600_ of 0.6. ER derived membranes were purified by differential centrifugation and immuno-isolation and subsequently analyzed by quantitative shotgun lipidomics. **(A)** Lipid class composition given as mol% of all lipids in the sample. Sterol (Erg, ergosterol), storage lipids (EE, ergosteryl ester; TAG, triacylglycerol) and membrane glycerolipids (DAG, diacylglycerol; PA, phosphatidic acid; PC, phosphatidylcholine; PE, phosphatidylethanolamine; PI, phosphatidylinositol; PS, phosphatidylserine). **(B)** Rare membrane glycerolipids (CDP-DAG, cytidine diphosphate diacylglaycerol; PG, phosphatidylglycerol; CL, cardiolipin), lysolipids (LPC, lyso-phosphatidylcholine; LPE, lyso-phosphatidylethanolamine; LPI, lyso-phosphatidylinositol) and sphingolipids (Cer, ceramide; IPC, inositolphosphorylceramide; MIPC, mannosyl-IPC; M(IP)_2_C, mannosyl-di-IPC). **(C)** Lipid metabolic pathway of PI biogenesis. **(D)** Total number of double bonds in membrane glycerolipids (except CL which has four acyl chains) as mol% of this category. **(E)** Change of average acyl chain length and unsaturation upon inositol depletion. Dot diameters are proportional to abundance of the respective lipid class in the ER membrane (as in Figure 3A) of indicated growth condition. Statistical significance was tested by multiple t tests correcting for multiple comparisons using the method of Benjamini, Krieger and Yekutieli, with Q = 1 %, without assuming consistent standard deviations. *p < 0.05, **p < 0.01, ***p < 0.001.

We found that inositol-depletion causes a substantial reduction of inositol-containing PI lipids (Figure 3A, B, Supplementary Table S3). This is accompanied by a drastic accumulation of CDP-DAG lipids, which serve as direct precursors for PI synthesis via Pis1 (Henry *et al*, 2012) (Figure 3A, B). Even the penultimate precursor of PI synthesis, PA, is found at significantly increased levels in the ER upon inositol-depletion (Supplementary Figure S3A, D). Sphingolipids, whose hydrophilic lipid headgroups also contain inositol, are not depleted under this condition (Figure 3B). This implies a strict prioritization for sphingolipid biosynthesis over PI synthesis when inositol becomes limiting. Overall, the molecular lipid fingerprint of the lipid bilayer stress caused by inositol-depletion is characterized by substantial changes in the abundance of anionic lipids, PI in particular (Figure 3A, B).

We further dissected the compositional changes of the ER membrane lipidome upon inositol-depletion at the level of the lipid acyl chains and observed a global trend toward shorter and more saturated glycerophospholipids (Figure 3D, E). While these changes are likely to fine-tune the physicochemical properties of the ER membrane, it is unlikely that these changes alone are sufficient to trigger the UPR by activating Ire1 (Halbleib *et al*, 2017). Hence, it is tempting to speculate that the overall reduction of anionic lipids, which directly affects the negative surface charge density of the ER (Supplementary Figure S4A, G), contributes to lipid bilayer stress. While a mechanistic role of individual lipids and collective membrane properties on UPR activation can be established only *in vitro*, our data provide a quantitative basis for studying the contribution of lipids and bulk membrane properties to chronic ER stress after a biochemical reconstitution of UPR transducers in native-like membrane environments.

### Lipid bilayer stress caused by proteotoxic agents Tunicamycin (TM) and Dithiothreitol (DTT)

Prolonged proteotoxic stresses activate the UPR via a membrane-based mechanism (Promlek *et al*, 2011; Väth *et al*, 2021) but the molecular underpinnings remain unknown. To address this gap, we exposed exponentially growing cells for 4 h to either 2 mM DTT or 1.5 μg/ml TM in SCD medium, isolated ER-derived membranes via MemPrep, and performed comprehensive lipidomic (Figure 4) and proteomic analyses (Figure 5). Our goal was establishing ER membrane compositions, which are known to trigger the UPR, rather than studying the impact of proteotoxic stress and UPR signaling on the membrane composition.

**Figure 4.**
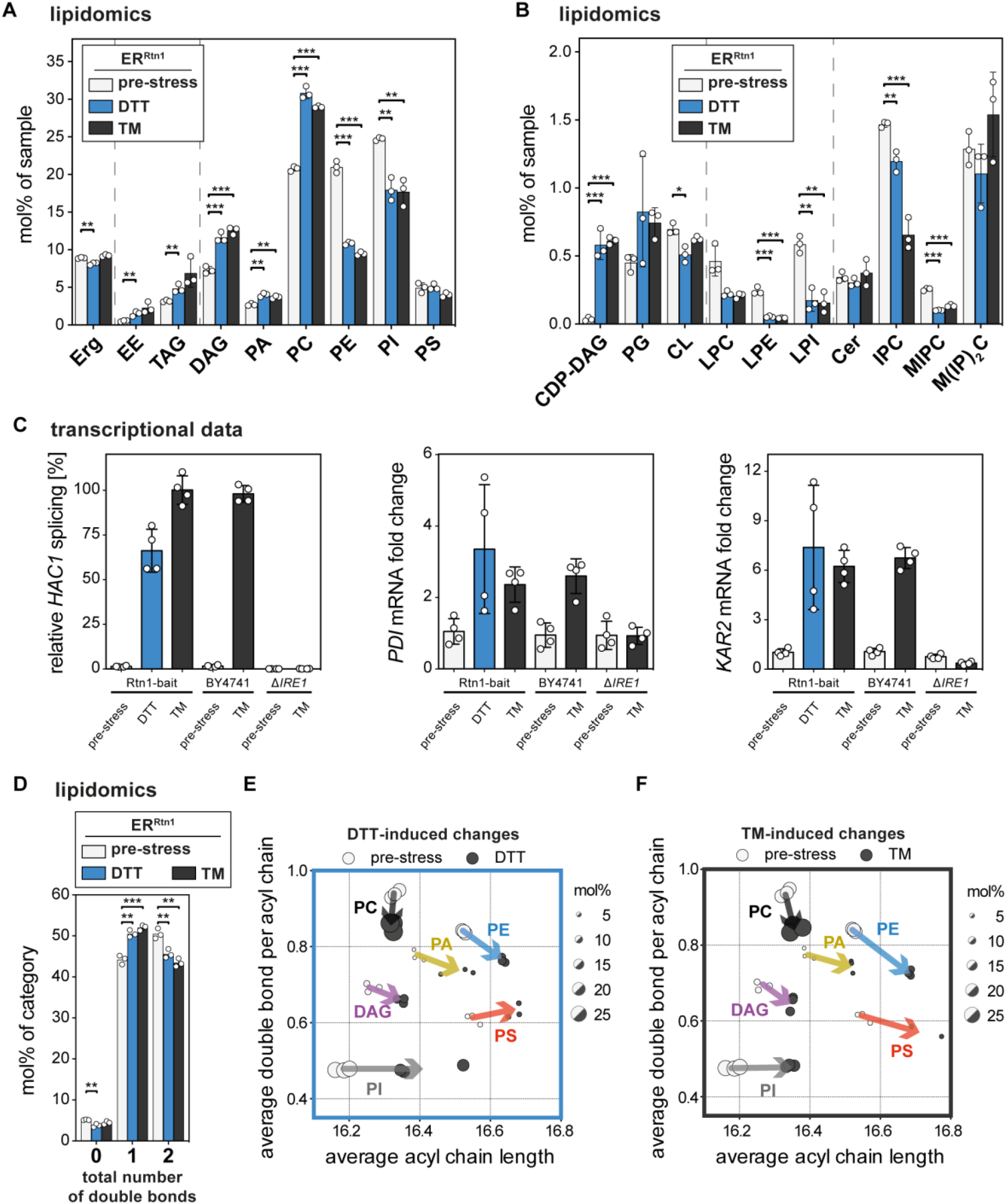
ER lipidomes of DTT- and TM-stressed cells indicate a shift towards a thicker, more saturated membrane. SCD_complete_ medium was inoculated with Rtn1-bait cells to an OD_600_ of 0.1 from an overnight pre-culture. Cells were grown to an OD_600_ of 0.8 and then stressed by addition of either 2 mM DTT or 1.5 μg/ml TM for 4 h. ER derived membranes were purified by differential centrifugation and immuno-isolation and subsequently analyzed by quantitative shotgun lipid mass spectrometry. **(A)** Lipid class distribution of sterol, storage lipids and abundant membrane glycerolipids in ER-derived vesicles from cells that were either challenged with 2 mM dithiothreitol (DTT) or 1.5 μg/ml TM for 4 h. The ER lipidome undergoes significant remodeling upon ER stress. **(B)** Lipid class distribution of rare membrane glycerolipids, lysolipids and sphingolipids. **(C)** Cells were grown as described above. UPR activation was measured by determining the levels of spliced *HAC1* mRNA and the mRNA of downstream UPR target genes (*PDI, KAR2*) before and after 4 h of DTT or TM treatment. Data for relative *HAC1* splicing was normalized to the TM treated Rtn1-bait condition. *PDI* and *KAR2* mRNA fold changes were calculated as 2^-ΔΔCT^ and normalized to Rtn1-bait pre-stress. **(D)** Total number of double bonds in membrane glycerolipids (without CL) given as mol% of this category. **(E)** Changes in average acyl chain length and unsaturation of main glycerolipid classes upon DTT induced ER stress. Dot diameters are proportional to abundance of the respective lipid class in the ER membrane (as in Figure 4A) of indicated growth condition. **(F)** Changes in average acyl chain length and unsaturation of main glycerolipid classes upon TM induced ER stress. Statistical significance was tested by multiple t tests correcting for multiple comparisons using the method of Benjamini, Krieger and Yekutieli, with Q = 1 %, without assuming consistent standard deviations. *p < 0.05, **p < 0.01, ***p < 0.001.

**Figure 5.**
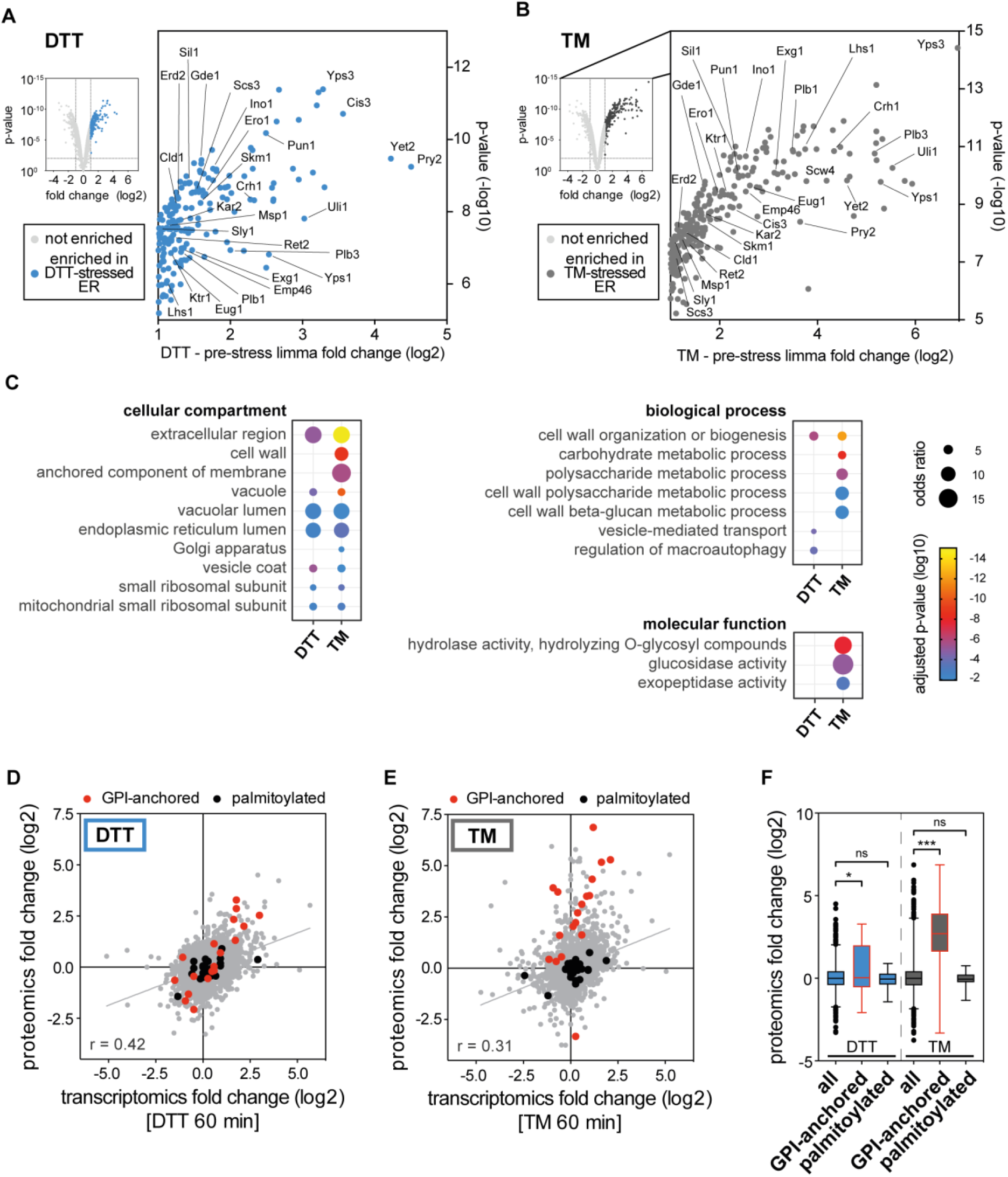
The proteome of the ER under conditions of prolonged proteotoxic stress. ER-derived vesicles were isolated by MemPrep and subsequently analyzed by untargeted proteomics. An additional sodium carbonate wash step was performed on P100 to remove soluble proteins from the membrane preparation. **(A)** Limma analysis identified proteins that are accumulating in ER preparations after prolonged DTT-induced stress (blue dots, top right quadrant of volcano-plot). Proteins that are discussed in the text are indicated. **(B)** Limma analysis showing proteins that are accumulating in the ER upon prolonged TM-induced ER stress (gray dots, top right quadrant of volcano-plot). Proteins that are discussed in the text are indicated. Accumulation of proteins in stressed ER was considered significantly when they were at least enriched two-fold compared to pre-stress with a p-value <0.01. **(C)** Enriched gene ontology terms (GO terms) in the list of proteins that are accumulating in ER-derived vesicles under the indicated ER stress conditions. GO terms are grouped by categories, FDR <1 %. **(D)** Correlation of previously published transcriptome data after one hour of DTT stress with our proteomics data after 4 h DTT-induced ER stress (Pearson correlation r = 0.42). GPI-anchored proteins (red dots) are clustering above the line of linear regression in contrast to palmitoylated proteins (black dots). **(E)** Correlation of previously published transcriptome data after one hour of TM stress with our proteomics data after 4 h TM-induced ER stress (Pearson correlation r = 0.31). In contrast to palmitoylated proteins (black dots) GPI-anchored proteins (red dots) are clustering above the line of linear regression. **(F)** Median limma fold changes over pre-stress condition of GPI-anchored, palmitoylated and all identified proteins. Whiskers indicate 1-99 percentile, significance was tested by Kolmogorov-Smirnov test. *p < 0.05, **p < 0.01, ***p < 0.001.

Surprisingly and despite each stress being completely different in its mechanism of action, the ER membranes isolated from either DTT-or TM-stressed cells have virtually identical lipid compositions, both profoundly distinct from the unstressed ER (Figure 4A, B, Supplementary Table S3). In fact, principal component analysis (PCA) shows clear separation between the stressed and unstressed ER, whilst highlighting the self-similarity of individual replicates (Supplementary Figure S4B). The two principal components PC1 and PC2 together defined >62 % of the lipidomic variation across all samples, with PC1 comprising 35.8 % of that variance. Notable segregation along PC1 was observed between the unstressed and the TM- and DTT-stressed conditions. With respect to the lipid composition, the stressed ER features higher levels of neutral storage lipids (EEs and TAGs) (Figure 4A), which may be caused by a reduced growth rate and an increased flux of fatty acids into storage lipids, as previously suggested, or a depletion of lipid metabolites from the medium (Listenberger *et al*, 2003; Vevea *et al*, 2015; Henne *et al*, 2018; Reinhard *et al*, 2020). The unusually high level of the lipid metabolic intermediates CDP-DAG and DAG in the stressed ER is consistent with these possibilities (Figure 4A, B). Notably, we confirmed that DTT and TM indeed trigger the UPR by RT-qPCR experiments (Figure 4C). These control experiments also show that the presence of the bait in the ER does not perturb UPR activation under these conditions (Figure 4C).

Compared to the PC-to-PE ratio of 1.0 in the unstressed ER, both the DTT- and the TM-stressed ER feature strikingly increased PC-to-PE ratios of 2.8 and 3.1, respectively, similar to what we noticed for the choline challenge (Figure 4A; Supplementary Figure S3A-C). Because the aberrantly high PC-to-PE ratio of ∼2.4 observed upon choline challenge (Supplementary Figure S3A) does not activate the UPR (Supplementary Figure S3F), while inositol-depletion does so even without perturbing the PC-to-PE ratio (Figure 3A, Supplementary Figure S3F), it is unlikely that increased PC-to-PE ratios observed under conditions of prolongd proteotoxic stresses can directly trigger the UPR (Fu *et al*, 2011; Gao *et al*, 2015; Ho *et al*, 2020; Ishiwata-Kimata *et al*, 2022). However, in both the DTT- and the TM-stressed ER, we find reduced levels of negatively charged, inositol-containing lipids (PI, LPI, IPC, MIPC) (Figure 4A, B), which is only partially compensated by mildly increased levels of PA and CDP-DAG (Figure 4A, B; Supplementary Figure S4A). Furthermore, we find that the glycerophospholipids of the stressed ER are significantly longer and more saturated compared to the unstressed ER (Figure 4D-F). Because Ire1 uses a hydrophobic mismatch-based mechanism (Halbleib *et al*, 2017), it is possible that these mild changes in the acyl chain region synergize with the reduction of anionic lipids in the ER membrane to mount a robust UPR. The molecular fingerprints of lipid bilayer stress therefore provide an important framework for dissecting the role of anionic lipids in UPR activation *in vitro*.

Based on the detailed molecular information, we established ER-like lipid compositions mimicking the stressed and unstressed ER using twelve commercially available lipids (Supplementary Table S2). The ER-like lipid mixtures were chosen to match for each condition the lipid class composition, the overall degree of lipid saturation, and the acyl chain composition in each individual lipid class. We performed all-atom molecular dynamics (MD) simulations on these ER-like compositions (Supplementary Figure S4C). Remarkably, all three ER-like lipid mixtures are substantially different to a lipid bilayer composed only of PC lipids with respect to membrane thickness (Supplementary Figure S4D), lipid packing defects (Supplementary Figure S4E), and the free volume profile (Supplementary Figure S4F). A particularly intriguing difference between the stressed and unstressed ER-like mixtures is the significantly different distribution of positive and negative charges in the water-membrane interface (Supplementary Figure S4G), which reflects the different abundance of anionic lipids in the stressed ER (Supplementary Figure S4A). Hence, beyond establishing fingerprints of the stressed ER, we provide a resource for studying protein-lipid and protein-membrane interactions, which will help studying the structure and function of membrane proteins in realistic, native-like membrane environments.

### Proteomic analysis of the DTT- and TM-stressed ER

Our lipidomic analysis uncovered huge differences between the stressed and the unstressed ER. To compare these with the proteomic changes we used MemPrep and quantitative proteomics to uncover proteomic changes in the stressed ER. Prior to subjecting microsomal membranes to the immuno-isolation procedure, we washed the microsomes with sodium carbonate to remove loosely attached peripheral proteins and contaminating cytosolic proteins. A total of 2952 proteins were robustly detected in three biological replicates of both the stressed and unstressed ER. Globally, the ER proteomes of DTT- and TM-stressed cells are largely similar (Pearson correlation coefficient r = 0.82, Supplementary Figure S5A). We find that prolonged proteotoxic stress is associated with a substantial remodeling of the ER proteome and the accumulation of 1) UPR target proteins, 2) lipid metabolic enzymes, 3) membrane trafficking machineries including cargoes, as well as 4) cell wall proteins and cell wall biogenesis factors. Notably, the accumulation of proteins in the stressed ER can be due to a transcriptional upregulation via the UPR or due to a mislocalization of non-ER proteins to the ER.

UPR targets found accumulated in the stressed ER include ER-luminal (co-)chaperones such as Kar2, Sil1, and Lhs1 as well as proteins involved in disulfide bridge formation such as Eug1 and Ero1 (Figure 5A, B). Most profoundly accumulated is Uli1, a known UPR target of unknown function, and Yet2, a homolog of the mammalian BAP31, which has been implicated in ERAD (Wakana *et al*, 2008) and the formation of ER-mitochondria contacts (Namba, 2019).

Prolonged proteotoxic stress activates the UPR via a membrane-based mechanism (Promlek *et al*, 2011) and it is associated with the accumulation of various lipid metabolic enzymes in the ER thereby affecting sterols (e.g. Pry2 and Skm1) and the metabolism of PI and PC (e.g. Ino1 and Gde1) (Figure 5A, B). Also the phospholipases Plb1 and Plb3, which are crucial for lipid fatty acyl chain remodeling, accumulate in the ER (Figure 5A, B) (Renne *et al*, 2015). The accumulation of the mitochondrial phospholipase Cld1 (active towards CL) in the ER, suggests that protein sorting and mitochondrial import are disrupted by proteotoxic stresses. Also, the fatty acyl-coenzyme A (CoA) diphosphatase Scs3 accumulates in the stressed ER. This homolog of the mammalian FIT2 is crucial for maintaining ER structure during stress by enabling a normal storage of neutral lipids in lipid droplets (Yap *et al*, 2020; Becuwe *et al*, 2020). The grossly perturbed abundance of various lipid metabolic enzymes in the stressed ER is likely to contribute to the lipidomic changes observed for the stressed ER (Figure 4) and may, at least in part, reflect homeostatic responses to maintain ER membrane function upon stress.

Aberrant protein folding in the ER prevents the ER exit of various secretory and membrane proteins with major consequences on the entire secretory pathway (Travers *et al*, 2000; Jonikas *et al*, 2009). In fact, crucial components of the COP-II (e.g. Emp46, Sly1, and Plb3) and COP-I (Ret2) pathways accumulate in the stressed ER. Likewise, the HDEL receptor Erd2 (Semenza *et al*, 1990) and the mannosyl-transferase Ktr1 involved in N- and O-linked glycosylations accumulate in the stressed ER. It is tempting to speculate that the aberrant ER localization of the membrane trafficking machinery contributes to the membrane-based activation of the UPR under conditions of prolonged proteotoxic stress.

Particularly striking for the stressed ER is the substantial accumulation of cell wall components, cell wall biogenesis factors (e.g. Cis3, Pun1, Crh1, and Exg1), and GPI-anchored proteins (e.g. Yps3 and Yps1).

### DTT and TM have similar yet distinct impact of the ER proteome

To functionally annotate the complex proteomic changes, we determined the enrichment of gene ontology terms (GO terms) in all upregulated proteins (Figure 5C). Most enriched GO terms regarding cellular components are shared for both sample sets derived from the DTT- and TM-stressed ER (extracellular region, vacuole, vacuolar lumen, endoplasmic reticulum lumen, vesicle coat) consistent with a general block of secretion. However, this analysis also reveals qualitative differences between DTT- and TM-stressed ER. While DTT seems to act more prominently on vesicular transport and autophagic processes (regulation of macroautophagy) (Figure 5C), TM seems to affect more selectively hydrolytic enzymes and carbohydrate-related metabolic processes thereby leading to an aberrant ER accumulation of vacuolar proteins, cell wall, and GPI-anchored proteins. The UPR is a powerful stress response that controls the expression of a large variety of UPR target genes (Travers *et al*, 2000). We observe a robust correlation between the UPR-dependent transcriptional upregulation of UPR target genes as determined by Travers *et al*. and the protein level in the stressed ER (Figure 5D, E). The dramatic, selective accumulation of GPI-anchored proteins in the TM-stressed ER (Figure 5F), however, suggests that different types of proteotoxic stresses can have different proteomic fingerprints at the organelle level. Our data suggest that TM impedes ER exit of GPI-APs by interfering with GPI anchor remodeling (Fujita *et al*, 2011; Rodriguez-Gallardo *et al*, 2020) consistent with previous observations that defects in the maturation of GPI-Aps trigger the UPR via a membrane based mechanism (Jonikas *et al*, 2009; Promlek *et al*, 2011).

. To further investigate the differences of DTT- and TM-induced changes of ER proteomes, we performed K-means clustering of the proteomic data (Supplementary Figure S5B). The analysis of GO term enrichments for the individual clusters revealed a small group of proteins that were accumulated in the DTT-stressed ER but depleted in the ER from TM-stressed cells (Supplementary Figure S5B, C, cluster 2). These proteins are involved in copper and iron transport (Fre7, Ctr1, and Fre1), which is interesting because iron affects the clustering propensity of Ire1 and the amplitude of UPR signaling (Cohen *et al*, 2017). Proteins in cluster 1 and 6 show more pronounced accumulation for TM-induced stress. The GO terms enriched in these clusters are connected to the cell wall and cell wall-related carbohydrate metabolism, while cluster 6 shows particularly strong enrichments of the GO terms vacuolar lumen and peptidase activity for the TM-stressed ER.

Taken together, our proteomics data suggests that DTT- and TM-triggered ER stress leads to globally similar, yet qualitatively distinct forms of ER stress. Both forms of ER stress cause an accumulation of non-ER proteins in the ER, whose contribution to UPR activation remains to be systematically investigated.

### Demonstrating the broad applicability of MemPrep on vacuolar membranes

While MemPrep was optimized for the ER, we sought to make it widely applicable to any organelle. To this end, the general feasibility of MemPrep was demonstrated by isolating vacuolar membranes. Given that the vacuole receives membrane material from various sources via the secretory pathway, endocytosis, macroauthophagy, lipophagy, and direct lipid transfer, it was unclear what the lipid composition of the vacuole would be and which organelle it would resemble even though its lipid composition has been partially addressed previously (Schneiter *et al*, 1999; González Montoro *et al*, 2018). From our genome-wide bait library, we decided to use a strain that expresses a bait-tagged variant of Vph1, the abundant, ATP-driven proton pump in the vacuole that exposes its C-terminal end to the cytosol. With the only exception that we used more starting material than for the ER, we applied the same protocol for the subcellular fractionation (Supplementary Figure S6A) and immuno-isolation (Figure 6A). Immuno-blotting of the final isolate revealed the presence of two vacuolar membrane proteins (the Vph1-bait and the palmitoylated Vac8), while other organellar markers for the ER and light microsomes (Dpm1, Kar2, 40kDa), mitochondria (Por1), endosomes (Ypt7, Pep12), the plasma membrane (Pdr5, Gas1), and peroxisomes (Pcs60 and Pex14) remained undetectable (Figure 6A). This demonstrates the global utility of MemPrep to isolate organelles.

**Figure 6.**
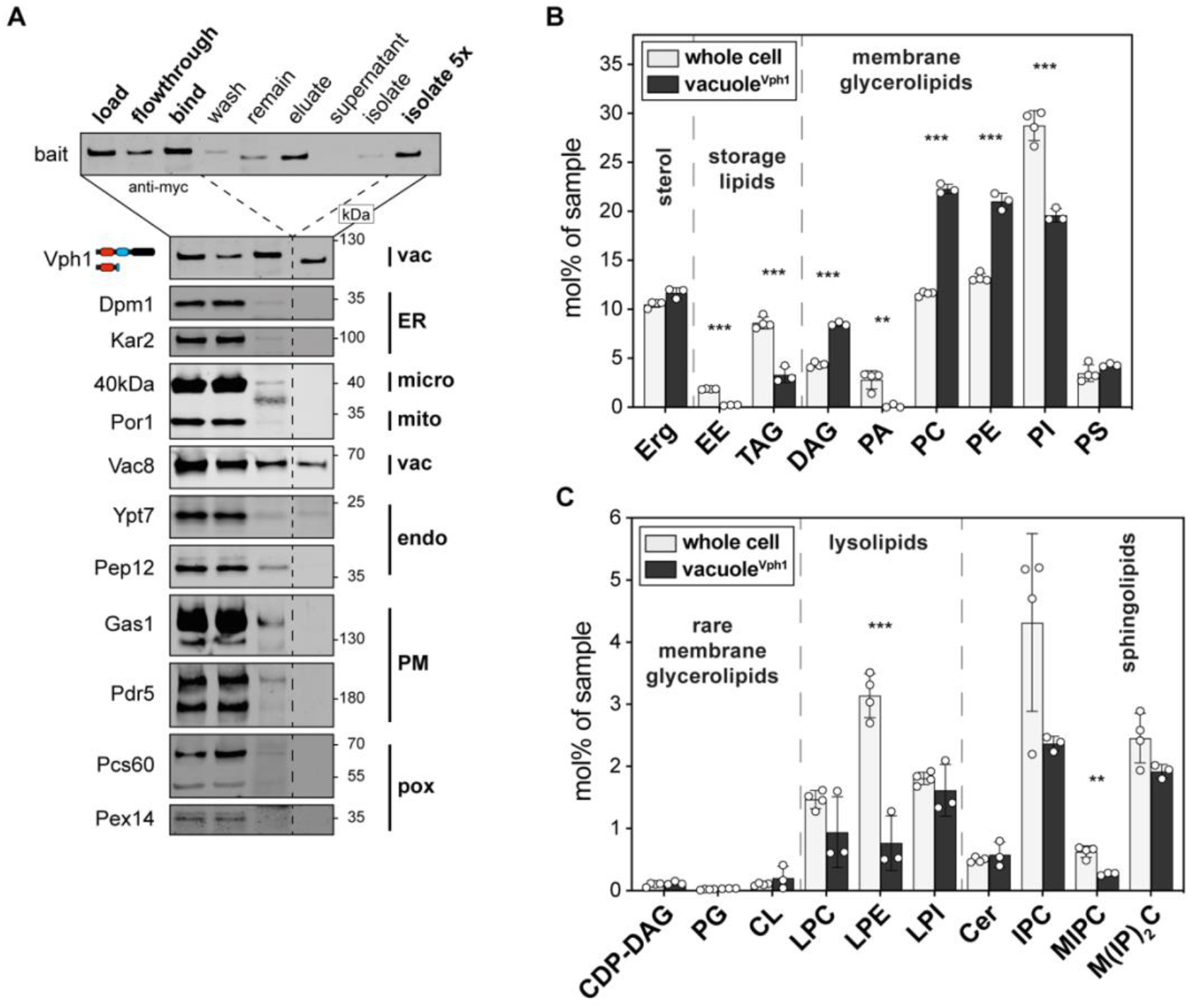
Lipid composition of the vacuolar membrane. **(A)** Immunoblot analysis of fractions after immuno-isolation via a vacuolar bait protein (Vph1-bait). Common organellar markers are shown: ER, endoplasmic reticulum (Dpm1 and Kar2); micro, microsomal fraction (40kDa); mito, mitochondria (Por1); vac, vacuole (Vac8); endo, endosomal system (Ypt7 and Pep12); PM, plasma membrane (Gas1 and Pdr5); pox, peroxisomes (Pcs60 and Pex14). 0.2 % of each fraction loaded per lane. **(B)** Lipid class composition given as mol% of all lipids in the sample. Classes are categorized into sterol (Erg, ergosterol), storage lipids (EE, ergosteryl ester; TAG, triacylglycerol), membrane glycerolipids (DAG, diacylglycerol; PA, phosphatidic acid; PC, phosphatidylcholine; PE, phosphatidylethanolamine; PI, phosphatidylinositol; PS, phosphatidylserine). **(C)** Continuation of lipid class composition given as mol% of all lipids in the sample. Classes are categorized into rare membrane glycerolipids (CDP-DAG, cytidine diphosphate diacylglycerol; PG, phosphatidylglycerol; CL, cardiolipin), lysolipids (LPC, lyso-phosphatidylcholine; LPE, lyso-phosphatidylethanolamine; LPI, lyso-phosphatidylinositol) and sphingolipids (Cer, ceramide; IPC, inositolphosphorylceramide; MIPC, mannosyl-IPC; M(IP)_2_C, mannosyl-di-IPC).Statistical significance was tested by multiple t tests correcting for multiple comparisons using the method of Benjamini, Krieger and Yekutieli, with Q = 1 %, without assuming consistent standard deviations. *p < 0.05, **p < 0.01, ***p < 0.001.

We then performed shotgun lipidomics on the isolated vacuolar membranes and found that the lipid composition is substantially different from the whole cell lysate with respect to neutral storage lipids, glycerophospholipids, and sphingolipids (Figure 6B, C, Supplementary Table S3). It is also remarkably distinct from the plasma membrane (Surma *et al*, 2011), while it features similar characteristics with the ER with respect to the abundance of PC, PI, and complex sphingolipids (Supplementary Figure S6B). Nevertheless, the vacuolar membrane is clearly distinct from ER by featuring significantly higher levels of ergosterol and DAGs. Most striking is the absence of PA lipids in the vacuolar membrane (Supplementary Figure S6B). Consistent with the vacuolar functions as lipid degrading organelle, we find higher levels of the lyso-lipids LPC, LPE, and LPI in the vacuolar membrane compared to the ER membrane (Supplementary Figure S6C). Furthermore, the lipid fatty acyl chains are more saturated in the vacuole compared to the ER membrane (Supplementary Figure S6D). These findings demonstrate the remarkably versatility of our immuno-isolation procedure and its feasibility for organellar lipidomics.

## Discussion

Understanding the homeostasis and adaptation of organellar membranes to metabolic perturbation and cellular stress is one of the key challenges in membrane biology. We developed MemPrep for the isolation of organellar membranes and a comprehensive and quantitative characterization of their composition. The versatility of this approach is demonstrated by the immuno-isolation of membrane vesicles from two very different organelles in yeast: the ER and the vacuole. Using state-of-the art lipidomics we provide a quantitative, molecular description of their membrane composition and establish a baseline for dissecting the role of lipids in transmembrane protein folding, trafficking, and function. Atomistic MD simulations highlight the difference between ER-mimetic membranes and PC-based lipid bilayers with respect to membrane thickness, lipid packing, the free volume profile, and surface charge distribution (Supplementary Figure S4D-G). The biochemical reconstitution of ER proteins in more realistic membrane environments is now feasible (Supplementary Table S2) and will become particularly relevant for the characterization of physicochemical membrane property sensors and the machineries that insert and extract membrane proteins into and out of the ER, respectively (Covino *et al*, 2018; Wu & Rapoport, 2021).

MemPrep overcomes the challenges associated with extensive membrane contact sites for the isolation of highly enriched organellar membranes. In contrast to recent strategies optimized for a rapid precipitation of organelles from yeast and mammalian cells (Liao *et al*, 2018; Melero *et al*, 2018; Ray *et al*, 2020; Higuchi-Sanabria *et al*, 2020), MemPrep maximizes for purity and provides access to the eluted membranes vesicles for a direct spectroscopic characterization and straightforward coupling to quantitative, analytical platforms. MemPrep provides a median enrichment of 25.7 for over 178 tested ER-resident proteins. This is remarkable, because even enrichments of 6 to 7 over the cell lysate have been considered as sufficient or even optimal in the past (Zinser & Daum, 1995).

Quantitative lipidomics show that ER lipids have a remarkably high content of unsaturated fatty acyl chains (75 mol%). The resulting low degree of lipid saturation is a crucial determinant of ER identity (Bigay & Antonny, 2012; Holthuis & Menon, 2014). It is continuously monitored by lipid saturation sensors (Covino *et al*, 2016; Ballweg *et al*, 2020), and actively maintained by the OLE pathway that controls the production of unsaturated fatty acids (Hoppe *et al*, 2000). The high fraction of unsaturated lipids in the ER may be crucial for the integration of transmembrane proteins in the ER membrane. Because transmembrane proteins differ substantially in their shape and hydrophobic thickness depending on their final subcellular destination and function (Sharpe *et al*, 2010; Quiroga *et al*, 2013; Lorent *et al*, 2017, 2020), a particularly soft and deformable membrane environment can provide a suitable environment for their folding and assembly in macromolecular complexes (Radanović & Ernst, 2021) and may also reduce the barrier for membrane protein integration via molecular invertases (Wu & Rapoport, 2021). In fact, the machineries that insert and remove membrane proteins into and from the ER, respectively, induce a local thinning of the membrane, presumably to lower the energy barrier for insertion and extraction (Wu & Rapoport, 2021). This membrane distortion should also render them sensitive to the lipid composition. In fact, increased membrane stiffness from increased lipid saturation or aberrantly high sterol levels inhibits the insertion of transmembrane proteins in model membranes (Brambillasca *et al*, 2005), the mammalian ER (Nilsson *et al*, 2001), and bacterial membranes (Kamel *et al*, 2022). Therefore, it comes as no surprise that the stiffness and thickness of the ER membrane is continuously monitored by the UPR for regulating the relative rate of protein and lipid biosynthesis as well as the ERAD machinery (Travers *et al*, 2000; Schuck *et al*, 2009; Halbleib *et al*, 2017).

The high level of DAG and unsaturated lipids may be further important for forming connections with other organelles via stalk-like structures and support lipid exchange. Indeed, recent simulations showed that polyunsaturated lipids and, to an even higher degree, DAG may stabilize membrane stalks by tens of kilojoule per mole (Poojari *et al*, 2021).

The ergosterol level in the ER is 9.7 mol%, which is consistent with previous estimations (Zinser & Daum, 1995; Schneiter *et al*, 1999; Van Meer *et al*, 2008). However, it is also higher than the level of cholesterol in the ER of mammalian cells, which is tightly maintained at ∼5 mol% (Radhakrishnan *et al*, 2008). This discrepancy becomes less surprising when considering the different impact of ergosterol and cholesterol on collective, physicochemical membrane properties (Atkovska *et al*, 2018). Despite their overall structural similarity, 10 mol% cholesterol increases the order and bending rigidity of a POPC bilayer at 25°C roughly 2-fold more than 10 mol% ergosterol (Henriksen *et al*, 2004).

Our ER lipid data are fully consistent with a gradual increase of lipid saturation along the secretory pathway (Brügger *et al*, 2000; Van Meer *et al*, 2008) and complement previous work on the composition of the trans-Golgi network / endosomal (TGN/E) system, secretory vesicles, and the plasma membrane in yeast (Klemm *et al*, 2009; Surma *et al*, 2011). However, they are not consistent with a functionally relevant increase of ergosterol along the early secretory pathway, because the level in the ER (9.7 mol%) is barely distinct from that of the TGN/E system (9.8 mol%) (Klemm *et al*, 2009). Hence, our findings support that sterols are sorted and enriched predominantly at the level of the TGN (Klemm *et al*, 2009). The lack of a robust sterol gradient in the early secretory pathway has important implications for the sorting of transmembrane proteins (Sharpe *et al*, 2010; Herzig *et al*, 2012; Quiroga *et al*, 2013; Lorent *et al*, 2017) and is consistent with recent observations that favor diffusion barriers established by a local enrichment of sterols as the basis of protein sorting (Weigel *et al*, 2021). This would be reminiscent of the ceramide-based diffusion barriers for membrane proteins in the ER between mother and daughter cells (Clay *et al*, 2014; Megyeri *et al*, 2019). Related to this, the presence of complex sphingolipids in the ER may seem surprising at first as the biosynthesis of complex sphingolipids occurs in the Golgi complex (Van Meer *et al*, 2008). Our direct, quantitative data provide evidence that complex sphingolipids can reach the ER at substantial rate, where they can be degraded by the sphingolipid-selective phospholipase C Isc1 for producing ceramides as part of a salvage pathway for sphingolipids (Matmati & Hannun, 2008).

The lipid composition of the vacuole is vastly distinct from that of the plasma membrane (Surma *et al*, 2011) despite a substantial intake of membrane material via the endocytic route (Wendland *et al*, 1998). However, the vacuolar membrane is also distinct from that of the ER with both lipid saturation (71 mol% unsaturated lipid acyl chains) and the sterol content (11.7 mol%) being higher in the vacuole. It is possible that a tighter packing of lipids in the vacuolar membrane is required to lower membrane permeability thereby contributing to the vacuolar acidification, which is crucial for activity of the luminal hydrolytic enzymes.

The most prominent lipid feature of the vacuole, which distinguishes it from the ER and other organellar membranes, is the absence of PA lipids. This is intriguing, because PA lipids are important signaling lipids that regulate lipid biosynthesis by sensing the cytosolic pH, which in turn is crucially regulated by the vacuolar proton pump (Young *et al*, 2010; Hofbauer *et al*, 2018). The higher level of lysolipids observed in vacuolar versus ER membranes is consistent with the role of the vacuole as a lipid-degrading organelle (Henry *et al*, 2012). Notably, due to the large head-to-tail volume ratio, lysolipids exhibit large positive intrinsic curvature that would favor the formation of membrane defects (Ting *et al*, 2018). Hence, the high levels of the negatively curved DAG may be required to counter-balance undesired effects from lysolipids on membrane stability.

We have established and employed MemPrep to identify molecular fingerprints of lipid bilayer stress. While lipid metabolic changes of the ER membrane have been firmly associated with chronic ER stress (Hotamisligil, 2010), the underlying molecular changes remained largely unexplored. We show that distinct conditions of lipid bilayer stress, namely inositol-depletion and prolonged proteotoxic stresses, are associated with dramatic changes of the ER lipid composition (Figure 3, 4). A PCA analysis reveals that the lipid fingerprints of inositol-depletion of prolonged proteotoxic stresses are remarkably distinct (Supplementary Figure S4B). We even observe opposing changes in the fatty acyl chain region: inositol-depletion is associated with a shortening of the lipid acyl chains, while prolonged proteotoxic stress correlates with acyl chain lengthening (Figure 3E, 4E, 4F). Furthermore, our quantitative data address an important, open question on the role of the PC-to-PE ratio as driver of the UPR. The ratio of PC-to-PE lipids is one of the key determinants of the lateral pressure profile and the curvature stress in cellular membranes, thereby affecting the conformational dynamics of membrane proteins (Marsh, 1996; van den Brink-van der Laan *et al*, 2004; Phillips *et al*, 2009). An aberrantly increased PC-to-PE ratio in the ER was suggested to cause chronic ER stress in obese mice (Fu *et al*, 2011), but the general validity of this interpretation is controversially discussed (Gao *et al*, 2015). We employed MemPrep, quantitative lipidomics, and sensitive UPR assays to investigate this point in yeast. Prolonged proteotoxic stress is associated with a dramatically increased PC-to-PE ratio in the ER, which goes well beyond the range of physiological variation observed at different growth stages (Janssen *et al*, 2000; Klose *et al*, 2012; Casanovas *et al*, 2015; Tran *et al*, 2019). In contrast, artificially increasing the PC-to-PE ratio by supplementing choline to the medium is not sufficient to trigger the UPR (Supplementary Figure S3F). Inositol-depletion, on the other hand, activates the UPR without significantly perturbing the PC-to-PE ratio (Supplementary Figure S3F). Hence, we show that an aberrantly increased PC-to-PE ratio is not sufficient to mount a robust UPR in yeast. We favor the idea than a decreased PC-to-PE ratio and the accumulation of lipotoxic intermediates trigger the UPR in yeast by activating Ire1 either directly or indirectly (Ho *et al*, 2020; Ishiwata-Kimata *et al*, 2022). Hence, our quantitative analysis of the ER membrane composition of stressed and metabolically challenged cells provide important insights to tackle mechanistic questions related to the chronic activation of the UPR.

Common to all tested conditions of lipid bilayer stress is an increase of saturated lipids in the ER membrane (Figure 3D, 4D) and a decrease in anionic lipids (Supplementary Figure S4A). While changes in lipid saturation have been firmly implicated in the activation of the UPR in both yeast and mammalian cells (Pineau *et al*, 2009; Volmer *et al*, 2013; Halbleib *et al*, 2017; Piccolis *et al*, 2019), a general role of anionic lipids as attenuators of the UPR has never been explored. We consider it highly unlikely that mildly increased levels of saturated lipids in the ER alone are sufficient to mount a full-blown UPR during inositol-depletion and prolonged proteotoxic stresses. We suggest that anionic lipids such as PI, PA, PS, and complex sphingolipids act as attenuators of the UPR, such that lipid saturation and the negative surface charge density of the ER jointly control output of the UPR. Notably, the level of PI and other inositol-containing lipids changes substantially in different growth stages (Casanovas *et al*, 2015) and the availability of inositol is limiting for optimal growth of the commonly used strain BY4741 (Hanscho *et al*, 2012). Integrating information on the membrane composition and properties, either directly or indirectly, is crucial for Ire1 to orchestrate membrane biogenesis by balancing the production of proteins and lipids (Covino *et al*, 2018).

Our proteomic analysis demonstrates the accumulation of a variety of proteins in the stressed ER, which are explained by a transcriptional upregulation via the UPR and aberrant trafficking along the secretory pathway (Figure 5). Most striking is the accumulation of GPI-anchored proteins under conditions of prolonged proteotoxic stresses. As aberrant handling of GPI-anchored proteins can trigger the UPR by a membrane-based mechanism (Jonikas *et al*, 2009; Promlek *et al*, 2011), we hypothesize that a failure to remodel of GPI-APs in the stressed ER (Rodriguez-Gallardo *et al*, 2020) triggers a build-up of these abundant cargoes, thereby perturbing the physicochemical properties of ER membrane and triggering the UPR (Halbleib *et al*, 2017). The mechanistic basis of this model will have to be addressed in the future. Based on our quantitative lipidomic and proteomic data, we propose that increased lipid saturation, depletion of anionic lipids, and changes of the membrane proteome all activate the UPR synergistically.

Combining MemPrep with quantitative proteomics unlocks a toolbox to study membrane protein targeting, sorting, and turnover at a global scale but with organellar resolution. Fascinating examples of inter-organelle communication highlight the crosstalk of the ER with lipid droplets, mitochondria, peroxisomes, and the vacuole in dealing with ER stress and lipotoxicity (Listenberger *et al*, 2003; Piccolis *et al*, 2019; Liao *et al*, 2021; Garcia *et al*, 2021). A rapid exchange of lipids between organelles provides a means to adapt to cellular stress and metabolic cues (Scorrano *et al*, 2019; Labbé *et al*, 2021). Recently developed approaches provide a first glimpse at the rate of lipid exchange between individual certain organelles (John Peter *et al*, 2022), but there is a great need for new preparative and analytical tools to keep track of all lipids at all times. The combination of biosensors providing high spatial and temporal resolution with MemPrep, which provides quantitative and comprehensive snapshots of a given organelle at a certain time, surfaces as a promising approach to study membrane adaptatively in a holistic fashion. We make MemPrep accessible to the community and have generated a collection of strains that facilitates the isolation of any organellar membrane of interest as demonstrated for the vacuolar membrane.

## Supporting information

Supplementary Table S1. Gene markers to calculate organelle enrichments.

Supplementary Table S2. In vitro and in silico lipid mixtures resembling the stressed and unstressed ER.

Supplementary Table S3. Lipidomics data. All lipidomics data in this study.

Supplementary Table S4. Analysis of protein enrichments and depletion during MemPrep of the ER using quantitative proteomics.

Supplementary Table S5. Prolonged proteotoxic stress causes substantial changes in the ER proteome.

---

**Supplementary Figure S1.**
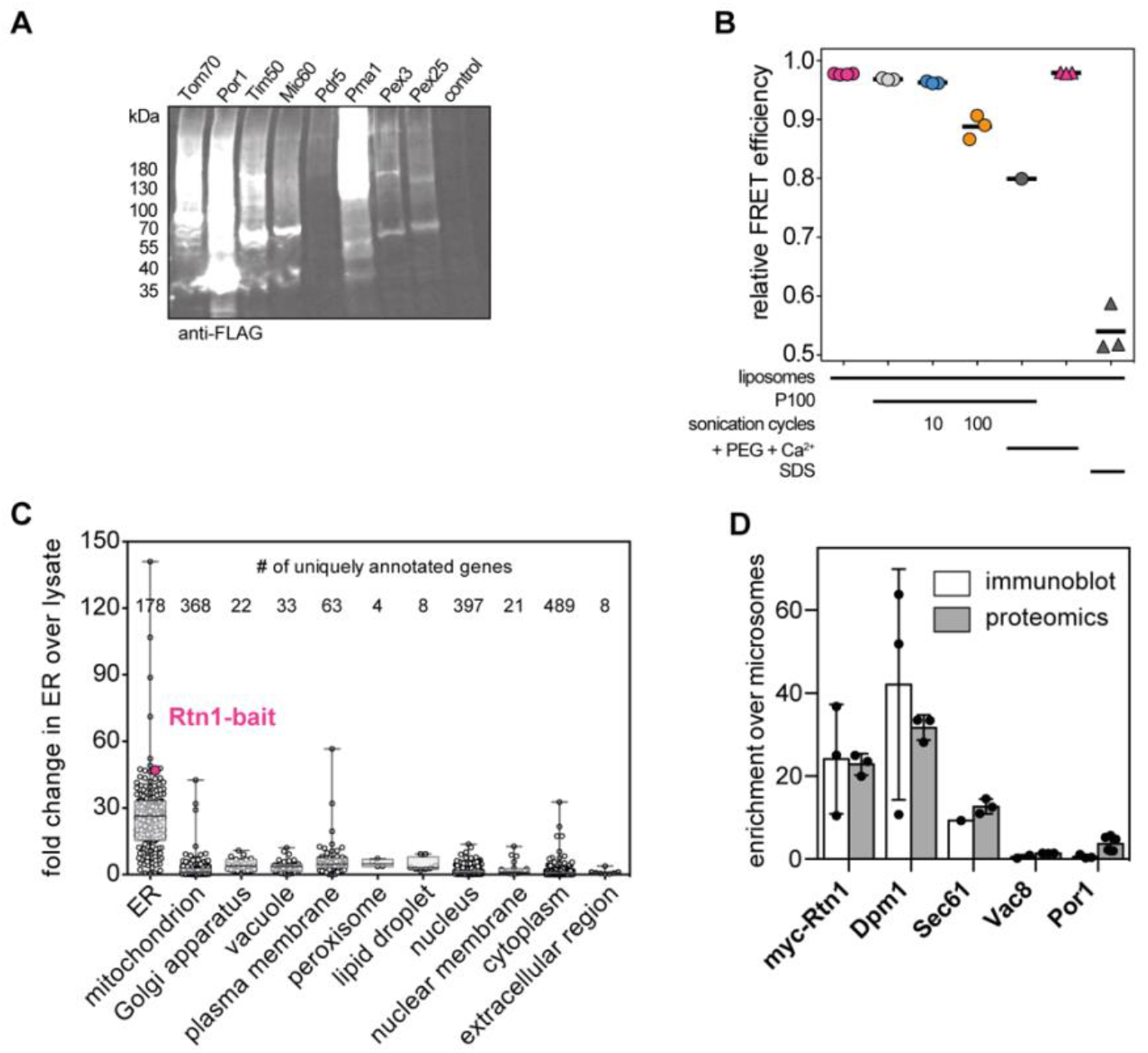
**(A)** From a systematic collection of strains in which every protein is tagged with a C-terminal bait tag (myc-3C-3xFLAG) we generated cell lysates for exemplary strains embedded in different organellar membranes. **(B)** Relative FRET efficiencies in mixtures of labeled liposomes and excess of unlabeled P100 microsomes after sonication, incubation with polyethylene glycol (PEG) and Ca^2+^, or sodium dodecyl sulfate (SDS). Lower relative FRET efficiency is the result of decreased average proximity of the two FRET-pair fluorophores and is therefore indicative for fusion of labeled liposomes with unlabeled P100 microsomes. **(C)** Number of genes with uniquely annotated cellular component gene ontology terms (indicated on the x-axis) that have been used to calculate organellar enrichments based on quantitative proteomics in Figure 1E. The fold change over the lysate of each individual potein in the ER fraction is plotted on the y-axis. The Rtn1-bait protein is highlighted in pink. Correlation of enrichments of organellar markers determined by either immunoblot analysis or proteomics. immuno-isolation bait protein (myc-Rtn1), ER markers Dpm1 and Sec61, vacuole marker Vac8, and outer mitochondrial membrane marker Por1.

**Supplementary Figure S2.**
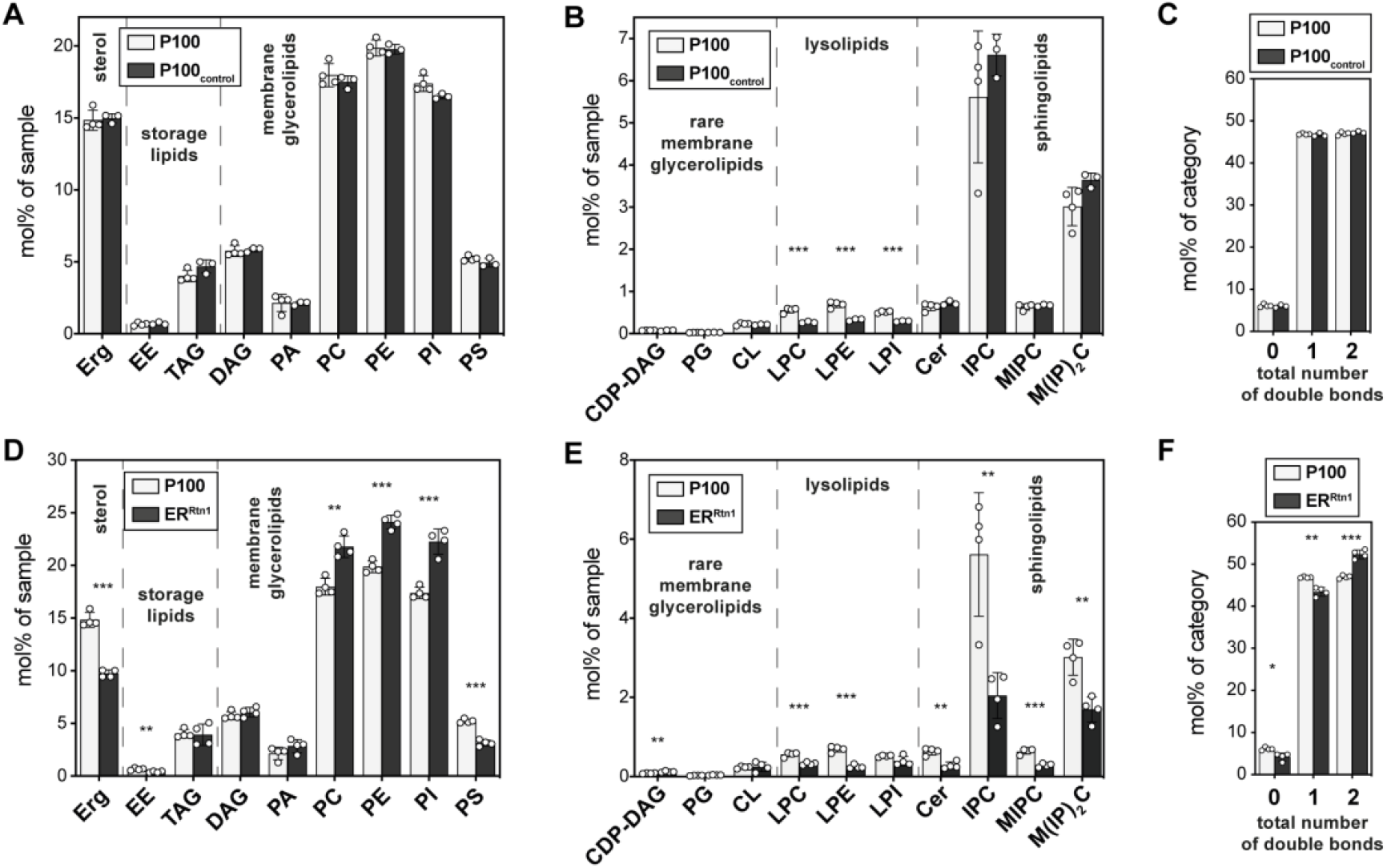
Lysolipids are depleted from the samples during the isolation procedure. To control stability of the sample an aliquot of P100 microsomes was incubated at 4 °C and overhead rotation (P100_control_) while the remaining sample was purified by immuno-isolation. **(A)** Abundance of detected lipid classes in microsomes (P100) and control microsomes after incubation for 8 h at 4 °C (P100_control_). **(B)** Lipid class distribution showing significantly less lyso-phospholipids in control microsomes (P100_control_). **(C)** The total number of double bonds in membrane glycerolipids is not changed. **(D), (E)** and **(F)** The lipid composition of the P100 crude membrane fraction before immuno-isolation (P100) is significantly different from the lipidome of ER vesicles derived by immuno-isolation via Rtn1 (ER^Rtn1^). Statistical significance was tested by multiple t tests correcting for multiple comparisons using the method of Benjamini, Krieger and Yekutieli, with Q = 1 %, without assuming consistent standard deviations. *p < 0.05, **p < 0.01, ***p < 0.001.

**Supplementary Figure S3.**
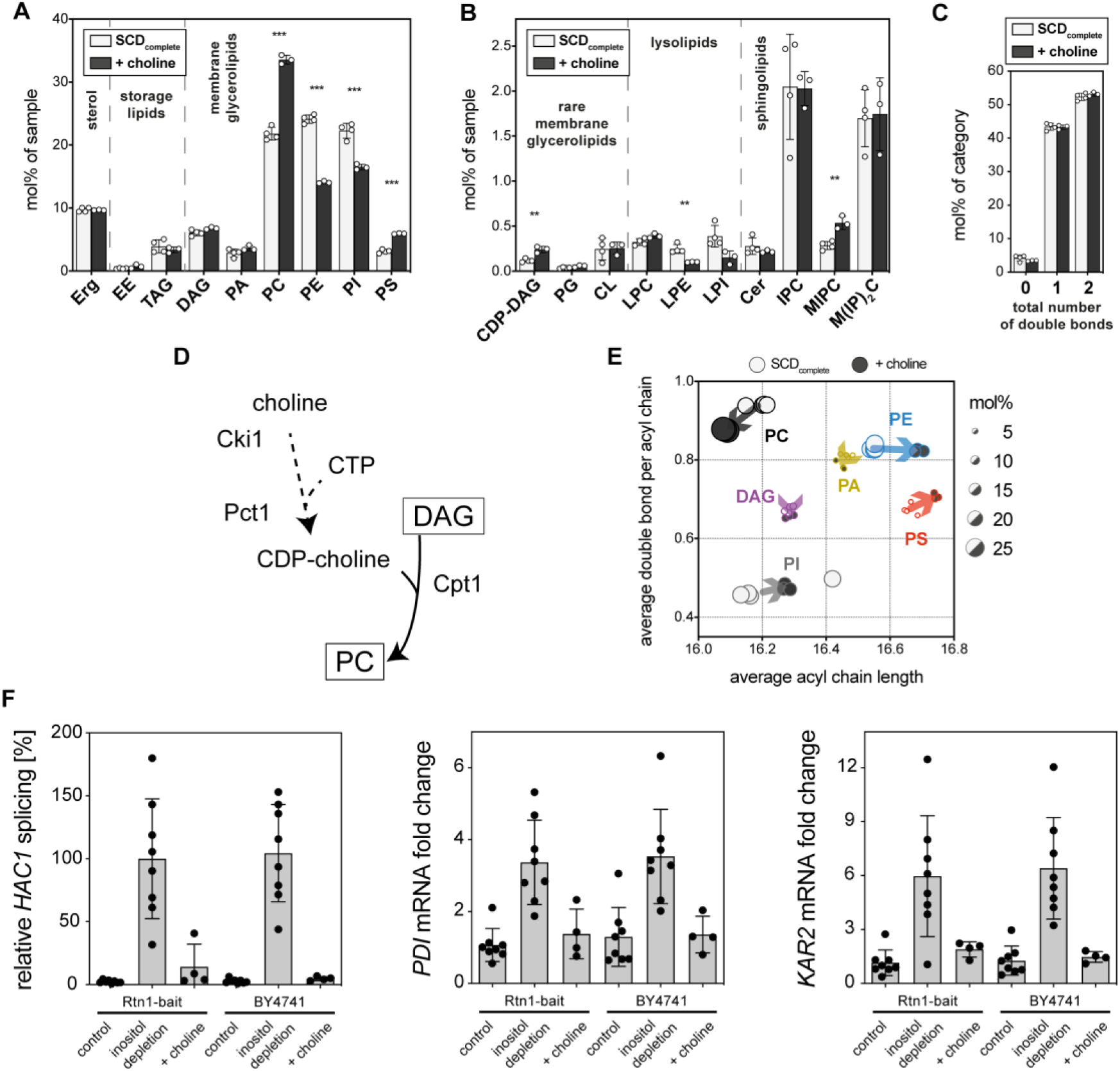
Metabolic interference with choline does not activate the UPR despite inducing dramatic lipidome changes. SCD_complete_ medium containing 2 mM choline (+choline) was inoculated with Rtn1-bait cells to an OD_600_ of 0.1 from an overnight pre-culture and cells were harvested at an OD_600_ of 1.0. ER derived membranes were purified by differential centrifugation and immuno-isolation and subsequently analyzed by quantitative shotgun lipidomics. **(A)** Lipid class composition given as mol% of all lipids in the sample. **(B)** Less abundant classes. **(C)** Total number of double bonds in membrane glycerolipids (except CL which has four acyl chains) as mol% of this category. Statistical significance was tested by multiple t tests correcting for multiple comparisons using the method of Benjamini, Krieger and Yekutieli, with Q = 1 %, without assuming consistent standard deviations. *p < 0.05, **p < 0.01, ***p < 0.001. **(D)** Lipid metabolic map of PC biosynthesis from external choline sources. **(E)** Changes in average acyl chain length and saturation of the main glycerophospholipid classes. Dot diameters are proportional to abundance of the respective lipid class in the ER membrane (as in Supplementary Figure S3A) of indicated growth condition. **(F)** Cells were grown as described above (+choline) or as described for inositol depletion experiments (Figure 3). In brief, Rtn1-bait cells with an OD_600_ of 1.2 were washed with inositol free medium and then cultivated for an additional 2 h in either inositol-free (inositol depletion) or SCD_complete_ medium (control). UPR activation was measured by determining the levels of spliced *HAC1* mRNA and mRNA of UPR target genes (*PDI* and *KAR2*). Data for relative *HAC1* splicing was normalized to Rtn1-bait cells under inositol depletion. *PDI* and *KAR2* mRNA fold changes were calculated as 2^-ΔΔCT^ and normalized to the Rtn1-bait control condition.

**Supplementary Figure S4.**
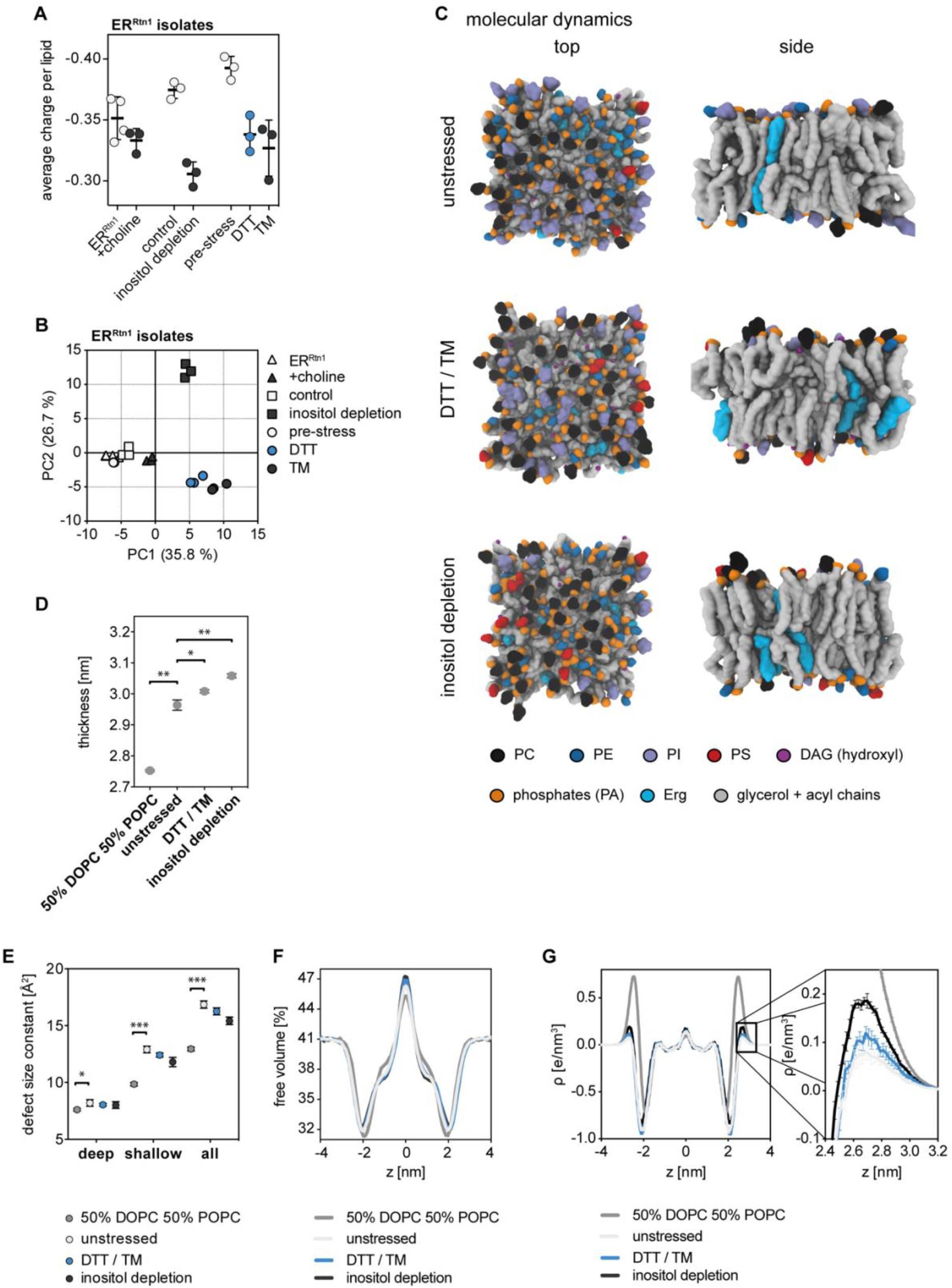
**(A)** Charge density of analyzed ER membranes represented as average charge per lipid. Net charges of the lipid classes were considered as follows: Erg 0, EE 0, TAG 0, DAG 0, PA -1, PC 0, PE 0, PI -1, PS -1, CDP-DAG -2, PG -1, CL -2, Cer 0, IPC -1, MIPC -1, M(IP)_2_C -2. **(B)** Principal component analysis (PCA) of lipidomics data from all ER-derived vesicle preparations. Based on 97 lipid molecular species that were detected in every sample. **(C)** Molecular dynamics (MD) simulations of proposed commercially available *in vitro* ER membrane lipid mixes for unstressed ER (unstressed) and ER under two different lipid bilayer stress conditions (DTT / TM, inositol depletion). Snapshots were taken after 100 ns. **(D)** Thickness measurements taken from MD simulations. **(E)** Determination of defect size constants in MD simulation of model membranes. **(F)** Free volume calculations from MD simulations. **(G)** The distribution of charges from MD simulations.

**Supplementary Figure S5.**
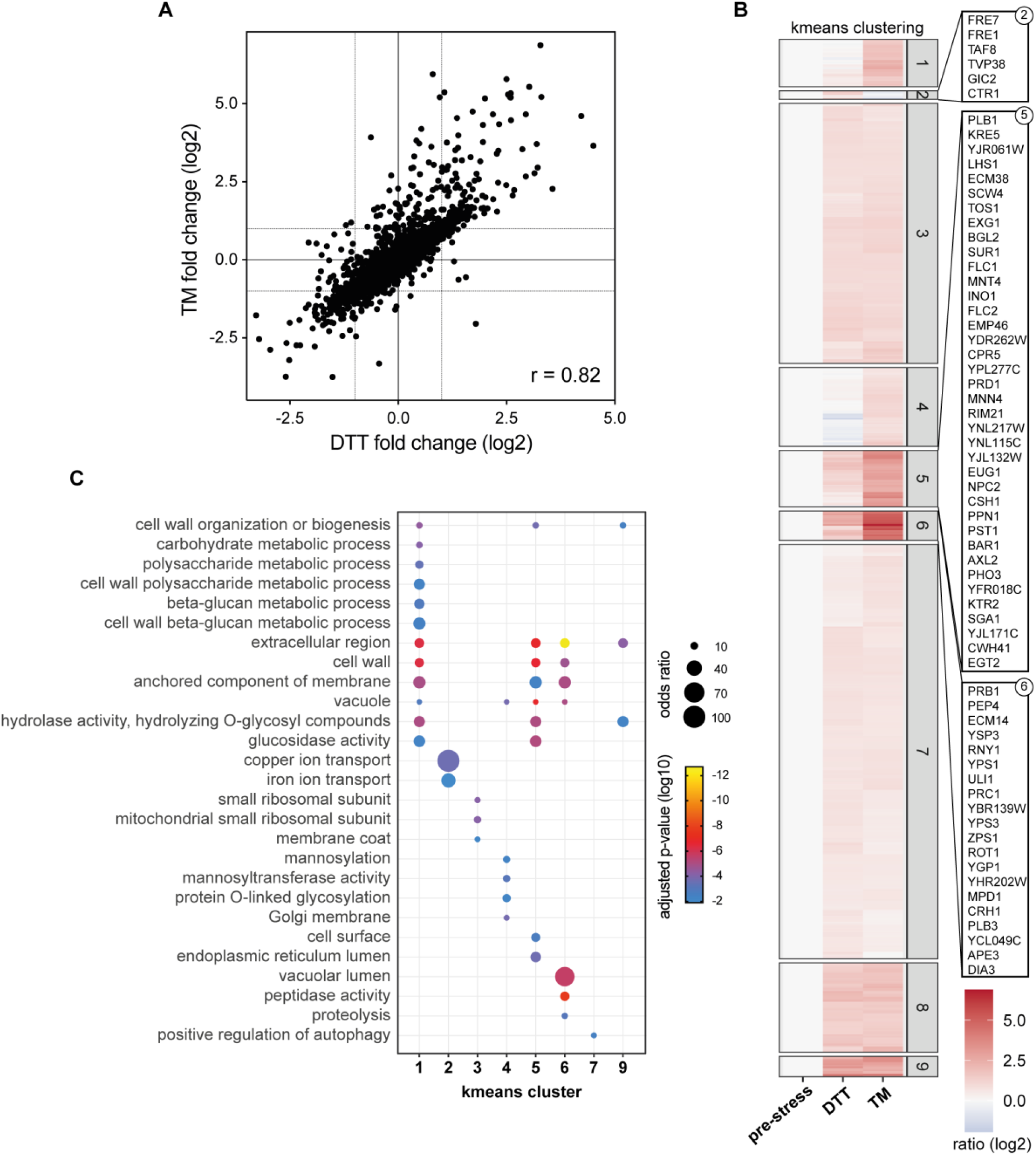
K-means clustering of DTT- and TM-induced changes in ER proteomes. **(A)** Correlation of DTT- and TM-induced limma fold changes over pre-stress with a Pearson correlation coefficient r = 0.82. **(B)** K-means clustering of proteins accumulating in the ER upon prolonged DTT-or TM-induced ER stress. **(C)** Gene ontology term enrichments in K-means clusters.

**Supplementary Figure S6.**
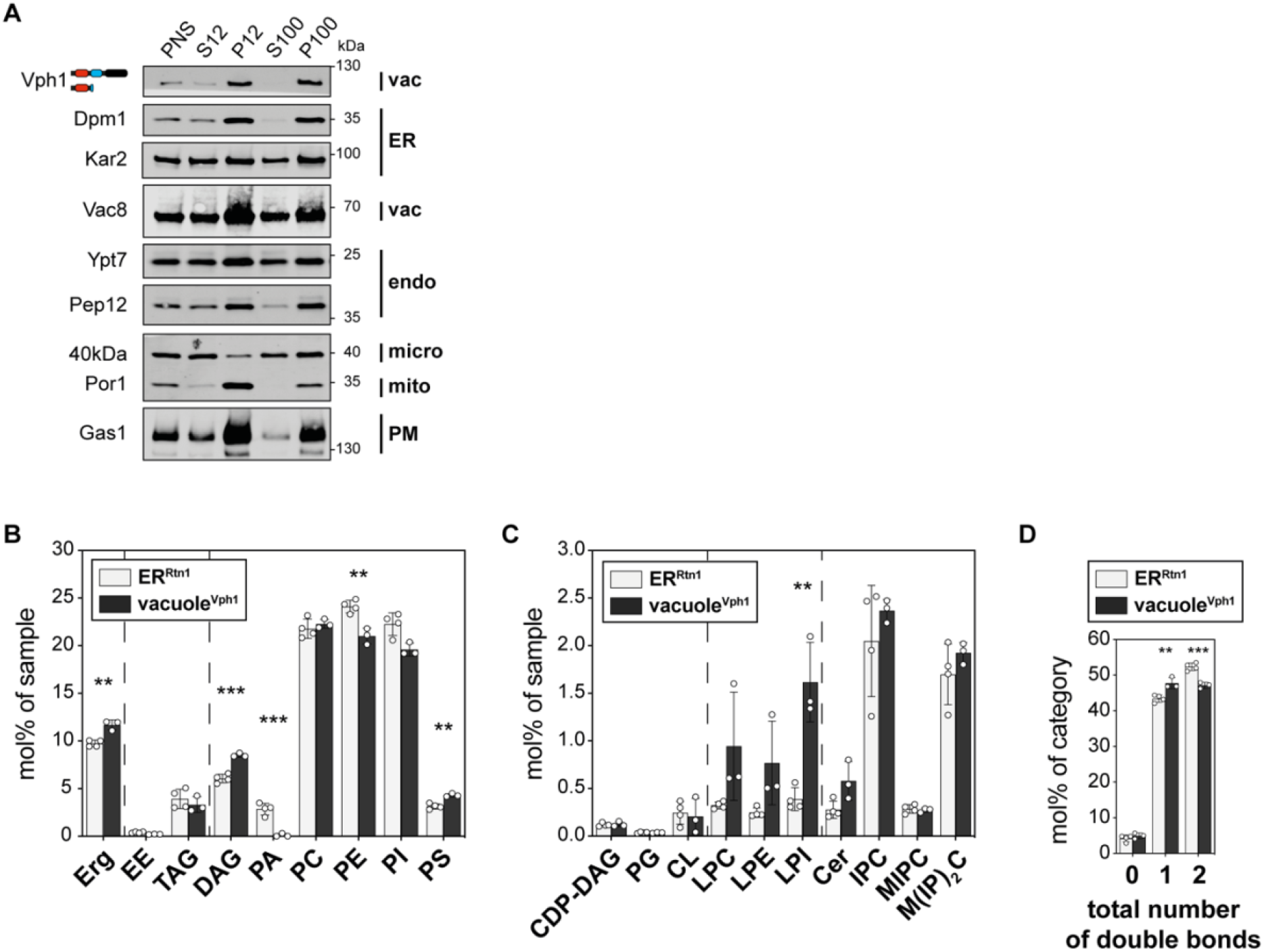
Lipidomics of the vacuole. **(A)** Differential centrifugation. **(B)** Lipid composition. **(C)** Continuation of lipid composition. **(D)** Saturation degree of membrane glycerolipids. Statistical significance was tested by multiple t tests correcting for multiple comparisons using the method of Benjamini, Krieger and Yekutieli, with Q = 1 %, without assuming consistent standard deviations. *p < 0.05, **p < 0.01, ***p < 0.001.

**Supplementary Table S1. Gene markers to calculate organell enrichments**. All genes with unique gene ontology term annotations in the category cellular component that were used to calculate ER enrichment.

**Supplementary Table S2. *In vitro* and *in silico* lipid mixtures**. Proposed ER-like lipid compositions for unstressed ER and different forms of lipid-bilayer stress (DTT / TM, inositol depletion) based on our lipidomics data. All lipids are commercially available to enable *in vitro* use.

**Supplementary Table S3. Lipidomics data**. All lipidomics data in this study.

**Supplementary Table S4. Analysis of protein enrichments and depletion during MemPrep of the ER using quantitative proteomics**. All proteomics data related to the validation of the ER isolation in Figure 1E.

**Supplementary Table S5. Prolonged proteotoxic stress causes substantial changes in the ER proteome**. All proteomics data related to the data presented in Figure 5.

## Acknowledgements

We wish to thank Michael Knop for generously providing the C’ SWAT library. We would like to thank Sepp Kohlwein, Karin Römisch, Christian Ungermann, Howard Riezman, Karl Kuchler, and Ralf Erdmann for providing antibodies and Sarah L. Keller, Georg Pabst as well as Chris Stefan for fruitful discussions and helpful comments. This work was funded by the VW foundation (Life?, #93089, #93092, #93090) to R.E., M.S., and J.S., by the Deutsche Forschungsgemeinschaft in the framework of the SFB894 to R.E. and the SFB1027 to J.H., and R.E., and by the European Research Council under the European Union’s Horizon 2020 research and innovation program (grant agreement no. 866011). MS is an incumbent of the Dr. Gilbert Omenn and Martha Darling Professorial Chair in Molecular Genetics.

