## Supplementary Table S1. Gene markers to calculate organelle enrichments. for "A new technology for isolating organellar membranes provides fingerprints of lipid bilayer stress"

ER (178) GO:0005783

AGP2, ALE1, ALG1, ALG12, ALG2, ALG3, ALG6, ALG7, ALG8, ALG9, ARE1, ARE2, ASI2, ATG15, ATX2, BIG1, BST1, CDC1, CDS1, CHO2, CNE1, CWH43, CYB5, DFG5, DFM1, DPL1, ELO1, ELO2, ELO3, EMC1, EMC4, EMC5, EMP46, EOS1, ERG11, ERG2, ERG24, ERG26, ERG28, ERG3, ERG4, ERG5, ERJ5, ERO1, ERV29, ERV41, ERV46, EUG1, FAR10, FAR3, FMN1, FRT1, GAA1, GAB1, GET2, GPI1, GPI10, GPI11, GPI13, GPI14, GPI16, GPI17, GPI2, GPI8, GTB1, GWT1, HLJ1, HMG2, HMX1, HNM1, HRD1, HRD3, HUT1, ICE2, IFA38, ILM1, INP54, IRC22, KAR1, KRE5, LAC1, LAG1, LAM5, LAS21, LCB1, LCB3, LIP1, MAM3, MNL1, MNL2, MNN10, MNN11, MNN9, MNS1, MPD2, MPS2, MSC7, NCR1, NNF2, NSG2, NTE1, OLE1, ORM2, OST1, OST2, OST3, OST6, OSW7, PBN1, PDI1, PER1, PER33, PEX31, PHO86, PKR1, PMT1, PMT2, PMT3, PMT4, PMT5, PMT6, PMT7, POM33, RFT1, SBH1, SCJ1, SCS3, SCS7, SCT1, SEC11, SEC20, SEC23, SEC39, SEC61, SEC62, SEC66, SEC72, SEI1, SFB3, SHR3, SIL1, SMP3, SOP4, SPC1, SPC2, SPC3, SRP101, SSH1, SSM4, SSS1, STE14, STE24, STT3, SVP26, TED1, TRS23, UBX7, UFE1, UIP4, USA1, USE1, USO1, VPH2, VPS62, VPS70, VTC2, VTC3, VTC4, WBP1, YET1, YET2, YET3, YOP1, YOS9, YPC1, YSY6, ZRG17, ZRT3

mitochondrion (368) GO:0005739

AAC3, AAT1, ABF2, ACK1, ACP1, AEP2, AFG3, AGC1, AIM10, AIM17, AIM18, AIM19, AIM23, AIM24, AIM36, AIM41, AIM45, AIM46, AIM9, ALD4, ALD5, ALO1, ALT1, ARC18, ARG5,6, ARG7, ARG8, ARH1, ATG9, ATM1, ATP11, ATP12, ATP16, ATP17, ATP18, ATP20, ATP3, ATP4, ATP5, ATP7, BAT1, BCH2, BIO2, BNA4, CAF4, CAT5, CBP2, CBP3, CBP4, CBP6, CCE1, CCM1, CCP1, CHD1, CIR1, CIR2, CMC1, COA1, COQ1, COQ11, COQ2, COQ3, COQ4, COQ5, COQ6, COQ8, COQ9, COR1, COX11, COX12, COX13, COX14, COX15, COX2, COX20, COX4, COX5B, COX6, COX9, CSF1, CTP1, CYC1, CYC2, CYC3, CYM1, CYT2, DIC1, DLD1, DLD2, DSS1, ECM31, EHD3, ERV1, ETR1, FAB1, FAU1, FMC1, FMP10, FMP25, FMP27, FMP32, FMP40, FMP41, FMP46, FMT1, FSF1, FYV4, FZO1, GCV1, GCV2, GCV3, GDH2, GEM1, GEP4, GEP7, GGC1, GLT1, GRX5, GUT2, HEM1, HEM14, HEM15, HEM25, HER2, HSP10, HTD2, IDH2, IDP1, ILV1, ILV2, ILV3, ILV5, ILV6, IMG1, IMG2, IMO32, INA22, INH1, IRA1, ISD11, KGD1, KGD2, KIP2, LAG2, LAM4, LAT1, LEU9, LIP2, LPD1, LSC1, LSC2, LYP1, LYS12, LYS4, MAM33, MAS1, MAS2, MBA1, MCP2, MCR1, MCT1, MCX1, MDL1, MDL2, MDM32, MDM38, MDV1, MGE1, MGM101, MGR1, MGR3, MIA40, MIC10, MIC12, MIC26, MIC27, MIC60, MIP1, MIR1, MMT1, MMT2, MNP1, MPM1, MRP1, MRP13, MRP20, MRP21, MRP4, MRP49, MRP51, MRP7, MRPL1, MRPL10, MRPL11, MRPL13, MRPL15, MRPL16, MRPL17, MRPL20, MRPL22, MRPL24, MRPL25, MRPL27, MRPL28, MRPL33, MRPL35, MRPL36, MRPL4, MRPL40, MRPL44, MRPL49, MRPL50, MRPL51, MRPL6, MRPL8, MRPL9, MRPS12, MRPS17, MRPS18, MRPS28, MRPS35, MRPS5, MRPS8, MRPS9, MRS1, MRS4, MRX1, MRX12, MSC6, MSD1, MSE1, MSH1, MSK1, MSM1, MSP1, MSS51, MST1, MTF2, MZM1, NAM9, NAT2, NCA2, NDE1, NDE2, NDI1, NFU1, NIT2, OAC1, OCT1, ODC1, ODC2, OM14, OM45, OMA1, OMS1, ORT1, OXA1, PAD1, PAM16, PAM17, PAM18, PCP1, PDA1, PDB1, PDX1, PET117, PET123, PET191, PET54, PET8, PET9, PHB1, PHB2, PIC2, PIM1, PNT1, POL1, POR2, POS5, PRP43, PSD1, PTC5, PTC7, PUT2, QCR2, QCR6, QCR7, QCR8, RCF1, RCF2, RDL2, RIM1, RIM2, RIP1, RMD9, RML2, RRF1, RRG1, RSM18, RSM19, RSM22, RSM23, RSM24, RSM25, RSM27, RSM28, RSM7, SAM35, SAM50, SCM4, SCO1, SCO2, SDH1, SDH2, SDH4, SDH5, SDH7, SDH8, SEN2, SEN34, SEN54, SHE9, SHM1, SHY1, SMC2, SOD2, SPT7, STE23, SUV3, SYT1, TAM41, TAO3, TAZ1, TCB3, TCD1, TCD2, TCM62, TFA1, TIM10, TIM11, TIM13, TIM17, TIM18, TIM21, TIM23, TIM44, TIM50, TIM9, TOM20, TOM22, TOM5, TOM70, TOM71, TRX3, TUF1, UBP16, UGO1, VAR1, YEA6, YHM2, YLH47, YMC1, YMC2, YME1, YME2, YML6, YMR31, YSC83

Golgi apparatus (22) GO:0005794

AKR1, APL5, APM3, APS3, BSC6, CCC2, GDA1, KRE2, KTR7, MNN1, MNN2, MNN5, MNT2, PEP12, RBD2, RUD3, SBE2, STV1, TMN3, VPS38, YIP5, YND1

vacuole (33) GO:0005773

APE3, ATG11, ATH1, AVT3, AVT4, AVT6, AXL2, CPS1, CSH1, CWH41, DCR2, ECM14, GNT1, HSE1, KEX1, KEX2, LCL2, MID2, NPC2, PRB1, RPP2B, SNA3, SRF1, STE13, TDA7, VAC14, VCX1, VMA8, VNX1, VPH1, VPS27, YCF1, YVC1

plasma membrane (63) GO:0005886

AIM44, ALR1, ARP3, BNI1, BSP1, BUD9, CAP1, CAP2, CCH1, CHS1, CLC1, END3, ENT2, ERG25, FCY2, FRE1, FTR1, FUN26, GIC2, GPR1, GSC2, HXT1, HXT3, HXT4, HXT5, IST2, KRE6, MNR2, MSC3, NRT1, PDR12, PHO87, PHO90, PIN2, PNS1, QDR2, QDR3, RGT2, SEC1, SEC15, SEG2, SKN1, SKT5, SLA2, SLM2, SLN1, SMF3, SNC1, SNC2, SSO1, SST2, SUL2, SYN8, TCP1, TPO1, TPO2, TPO3, TPO4, TRK1, VHT1, YAP1801, YEH2, YOR1

peroxisome (4) GO:0005777

ATG36, FOX2, PEX25, PEX27

lipid droplet (8) GO:0005811

ERG7, PET10, RRT8, TGL1, TGL3, TGL4, TGL5, YEH1

nucleus (397) GO:0005634

ABF1, ADA2, ADF1, AIR2, APC11, APM2, APN2, APS2, ARP4, ARP7, ARP9, ASG1, ASH1, BAS1, BDP1, BFR2, BMT2, BRE1, BRF1, BRR2, BRX1, BUD21, BUD22, BUR2, BUR6, CAF40, CDC45, CDC73, CEG1, CGI121, CGR1, CHA4, CHZ1, CIC1, CLP1, CSM1, CTF4, CTI6, CTR9, CWC15, CWC24, CYC8, DAL81, DBF2, DBP10, DBP3, DBP6, DBP7, DBP8, DBP9, DCC1, DEF1, DHR2, DIG1, DIG2, DIM1, DIP2, DLS1, DRS1, DST1, DUS1, EAF7, EBP2, ECM2, EFG1, ELF1, ENP2, ERB1, ERG13, ESF1, ETT1, FAL1, FCF1, FCF2, FHL1, FOB1, FPR3, FPR4, FYV6, FYV7, FZF1, GAR1, GCD10, GCD14, GCR2, GON7, GRC3, GTS1, GZF3, HAS1, HCA4, HHO1, HHT1, HIF1, HIR3, HMT1, HO, HPC2, HRQ1, HTZ1, IES2, IES3, IES6, IFH1, IMP3, IMP4, IOC4, IPI3, IRC5, ISW1, ISW2, ITC1, IXR1, JIP5, KAR3, KIP3, KRI1, KRR1, LCP5, LEO1, LEU3, LGE1, LHP1, LRP1, LRS4, LSM5, LYS14, MAC1, MAD1, MAG1, MAK16, MAK21, MAK5, MBP1, MCM10, MET1, MET4, MLP1, MMS1, MPH1, MPP10, MRC1, MSH2, MTR3, MTR4, MUS81, NAF1, NAN1, NCB2, NET1, NHP10, NHP2, NHP6A, NHP6B, NIS1, NKP1, NMA111, NMA2, NOC3, NOG2, NOP1, NOP10, NOP12, NOP13, NOP16, NOP2, NOP4, NOP53, NOP56, NOP58, NOP6, NOP7, NOP8, NOP9, NPL6, NRD1, NSA1, NSA2, NSE1, NSE4, NUG1, NUP192, NUT2, ORC1, PAF1, PAP1, PCT1, PDC2, PHO23, PIP2, PKH2, POL12, POL30, POL5, PPS1, PRP40, PRP45, PRP8, PSY2, PUF6, PUS1, PXR1, RAD1, RAD2, RAD33, RAV2, RCL1, RCM1, RCO1, RDH54, REB1, RET1, REX4, RFA2, RFC1, RFC2, RFC5, RIF1, RIX7, RKM4, RLF2, RLP24, RLP7, RNH202, RNH70, RNT1, ROK1, ROM2, ROX1, RPA190, RPA34, RPA49, RPB3, RPB5, RPB8, RPB9, RPC19, RPC25, RPC31, RPC40, RPC53, RPF1, RPF2, RPH1, RPI1, RPL28, RPL4A, RPL5, RPN13, RPN2, RPN7, RPO31, RPS13, RPS14B, RPS2, RPS3, RPS7A, RPS7B, RPS9B, RPT3, RPT4, RPT6, RRB1, RRN5, RRP1, RRP14, RRP15, RRP3, RRP36, RRP4, RRP40, RRP42, RRP45, RRP46, RRP5, RRP6, RRP7, RRP8, RRP9, RRS1, RRT14, RSA1, RSA3, RSA4, RSC3, RSC58, RSC6, RSC8, RSC9, RSE1, RTF1, RTT102, RTT106, RTT107, RTT109, RVB1, RVB2, RXT3, SAD1, SAS10, SCD5, SDA1, SEF1, SFH1, SFL1, SGD1, SGV1, SIF2, SKI3, SLX9, SMC1, SMC3, SMC5, SMI1, SMT3, SNF2, SNU114, SNU13, SNU66, SOF1, SOK1, SPB1, SPB4, SPN1, SPO14, SPT15, SPT8, SRN2, SRP14, SRP21, SRP40, SRP72, SRS2, SSF1, SSF2, SSN3, STE12, STH1, STO1, SUA7, SUB1, SUM1, SWC4, SWD3, SWI1, SYC1, TAF14, TAF7, TAF8, TBF1, TDA9, TFC1, TFC7, TFC8, THO1, THO2, THP1, THP3, TMA16, TMA23, TMC1, TOP1, TRA1, TRF5, TRM8, TUP1, UBC9, UBP6, UGA3, ULP1, URB1, URB2, URN1, UTP11, UTP13, UTP14, UTP15, UTP18, UTP21, UTP22, UTP25, UTP30, UTP4, UTP5, UTP6, UTP7, UTP8, UTP9, VMA22, WHI2, XRS2, YCS4, YGK3, YJU2, YPI1, YRA2, YTM1

nuclear membrane (21) GO:0031965

ALG14, ASM4, GTT3, NIC96, NSP1, NUP1, NUP100, NUP116, NUP120, NUP145, NUP157, NUP170, NUP188, NUP49, NUP53, NUP57, NUP84, NUP85, NUR1, PML39, SEH1

cytoplasm (489) GO:0005737

AAR2, ABP140, ABZ1, ABZ2, ACF2, ADD37, ADE12, ADE13, ADE16, ADE2, ADE4, ADE5,7, ADE6, AFG2, AGE1, AHA1, AIM2, AIM29, AKL1, ALD3, ALY1, ALY2, AMD1, ANT1, APL2, APM4, ARA1, ARA2, ARC1, ARC15, ARC35, ARG1, ARG4, ARK1, ARO1, ARO10, ARO2, ARO8, ARP6, ASK1, ASK10, ASN1, ASN2, ASP1, AST2, ATE1, ATG13, ATG19, ATG2, ATG20, ATG21, ATG26, ATG34, ATG8, ATS1, ATX1, AVL9, BBP1, BCK1, BDH1, BEM3, BET2, BOI2, BRE5, BRO1, BUD2, BUD27, BUG1, BUL1, BUL2, CAB3, CAF20, CCT2, CCT7, CCT8, CDC12, CDC15, CDC23, CDC27, CDC31, CDC37, CDC60, CEF1, CEX1, CHC1, CHS5, CIN2, CIN4, CKI1, CLU1, CMD1, CMK1, CMK2, CMP2, CNA1, CNM67, CNS1, COG2, COG4, COG5, COG8, CPA1, CPA2, CPR6, CPR7, CSE1, CSN12, CUE3, CUE5, CWC21, CWC22, CWC27, DAD3, DAK1, DAM1, DBF20, DCD1, DDR48, DED81, DEP1, DHH1, DID2, DID4, DMA1, DMA2, DOM34, DOS2, DPH1, DPH2, DPH5, DPH6, DRE2, DSS4, DTD1, DUO1, DYS1, ECM21, ECM25, ECM30, ECM32, ECM4, EDE1, EFB1, EFM2, EFM4, EFM5, EFT1, ELC1, ENT4, ENV7, ETP1, FAD1, FBP26, FDC1, FRA1, FRQ1, FRS1, FRS2, FUR1, GAD1, GCD1, GCD11, GCD2, GCD6, GCN3, GDE1, GDI1, GEA2, GET4, GGA1, GGA2, GIM3, GIM4, GIP4, GIR2, GIS2, GLC3, GLG2, GLO2, GLY1, GPH1, GPM2, GRX8, GSH1, GSH2, GUA1, GVP36, GYP6, GYP7, GYP8, HBS1, HCS1, HEL2, HIS1, HIS2, HIS4, HIS5, HIS7, HOM3, HSK3, HSP42, HUA2, ILS1, IMD2, IMD4, IMP2', INN1, INO1, INP51, INP52, INP53, IQG1, IRC24, IST1, ISY1, JNM1, JSN1, KEL1, KEL3, KES1, KIC1, KRS1, LAS17, LEA1, LEU1, LSB5, LSG1, LTE1, LYS2, LYS9, MAG2, MAK10, MCK1, MDE1, MDM20, MDR1, MEH1, MES1, MET10, MET12, MET14, MET2, MHP1, MHT1, MKK1, MKT1, MLF3, MOB1, MON1, MPT5, MRP8, MTW1, MUK1, MVD1, MYO2, NAB6, NAT3, NBA1, NIP1, NMD2, NMT1, NPA3, NRP1, NST1, NTF2, NTH1, NUD1, OCA1, OCA4, OLA1, OSH6, OSH7, OTU2, OXP1, PAA1, PAC10, PAM1, PAN2, PAN3, PAN5, PAR32, PBA1, PBS2, PBY1, PCS60, PDR17, PEP3, PEP5, PEP8, PEX22, PFD1, PFK26, PFY1, PGM1, PGM2, PHA2, PIG2, PKH1, PLP1, PLP2, PNP1, POL3, POP1, PPX1, PRK1, PRO1, PRO3, PRP28, PRS1, PRS3, PRS4, PRS5, PSA1, PTC4, PUF2, PUF3, PUF4, PXL1, PYC1, PYC2, RAM2, RAV1, RBA50, RBG1, RBG2, RBL2, RCK2, RCN2, REE1, REH1, REI1, RET2, RFS1, RGA1, RGA2, RGC1, RGD1, RIA1, RIB2, RIC1, RIM11, RMD1, RPD3, RPG1, RPL29, RPL30, RPN12, RPN5, RPP0, RPS1A, RPS1B, RQC1, RQC2, RTC5, RTG2, RTP1, RTT10, RVS161, RVS167, SAH1, SAM1, SAM2, SCD6, SCY1, SEA4, SEC2, SEC53, SEC7, SED4, SER1, SER3, SER33, SES1, SFM1, SFT2, SGN1, SGT2, SHE1, SHE3, SHE4, SIP5, SIW14, SKI2, SKI7, SKP2, SLF1, SLH1, SLK19, SLM4, SLP1, SMB1, SMY1, SMY2, SNU71, SNX3, SNX4, SNX41, SOL2, SPC19, SPC34, SPE1, SPE4, SQT1, SRP68, SSK1, SSK2, STE11, STE50, STE7, STI1, STP22, SUI2, SUP35, SUP45, SVL3, SWA2, TAE1, TAP42, TDA11, TDA2, TDA3, TEL2, TFS1, THI20, THI6, THI80, TIF5, TLG1, TMA108, TMA20, TMA22, TMA46, TMT1, TOA1, TOA2, TPS1, TPS3, TRL1, TRM11, TRM3, TRM44, TRM732, TRP2, TRP3, TRS120, TRS130, TSA1, TSL1, TSR4, TUB2, TVP23, TVP38, TWF1, TYR1, TYW3, UBA4, UBR1, UBR2, URA1, URE2, VAC8, VFA1, VHS2, VIP1, VMA2, VPS16, VPS17, VPS24, VPS28, VPS29, VPS3, VPS30, VPS35, VPS5, VPS9, WHI3, WHI4, WRS1, XKS1, XPT1, YAT2, YBP1, YBP2, YCG1, YHI9, YSC84, ZDS1, ZDS2, ZWF1

extracellular region (8) GO:0005576

BAR1, CRH1, DSE4, PHO5, SCW11, UTH1, UTR2, YGP1

ambiguous (1862)

AAH1, AAP1, AAT2, ABD1, ABP1, ACB1, ACC1, ACE2, ACF4, ACH1, ACL4, ACO1, ACO2, ACS2, ACT1, ADD66, ADE1, ADE17, ADE3, ADE8, ADH1, ADH3, ADH4, ADH5, ADH6, ADI1, ADK1, ADO1, ADP1, ADR1, AEP3, AFG1, AFI1, AFT1, AGE2, AGP1, AHP1, AIF1, AIM14, AIM20, AIM21, AIM25, AIM3, AIM39, AIM6, AIP1, AIR1, AKR2, ALA1, ALB1, ALD6, ALF1, ALG11, ALG13, ALG5, ALK1, ALK2, ALT2, AME1, AMN1, AMS1, ANP1, ANR2, AOS1, APA1, APC1, APC4, APD1, APE1, APE2, APE4, APJ1, APL1, APL3, APL4, APL6, APM1, APN1, APT1, AQR1, ARB1, ARC19, ARC40, ARD1, ARF3, ARG2, ARG81, ARI1, ARL1, ARL3, ARO3, ARO4, ARO7, ARO9, ARP2, ARP5, ARP8, ART5, ARX1, ASC1, ASF1, ASI1, ASI3, AST1, ATF1, ATF2, ATG18, ATG27, ATG40, ATG7, ATO3, ATP1, ATP10, ATP2, ATP23, AVO1, AVT1, AVT5, AVT7, AXL1, AYR1, AZF1, BAP2, BAP3, BAT2, BBC1, BCD1, BCH1, BCK2, BCP1, BCS1, BCY1, BDF1, BDF2, BDH2, BEM1, BEM2, BEM4, BER1, BET3, BET4, BFR1, BGL2, BIK1, BIM1, BIR1, BLM10, BMH1, BMH2, BMS1, BMT5, BMT6, BNA1, BNA3, BNA5, BNA6, BNA7, BNI4, BNI5, BNR1, BOI1, BOR1, BOS1, BPH1, BPL1, BPT1, BRE2, BRL1, BRN1, BRR1, BRR6, BSD2, BUB3, BUD13, BUD14, BUD17, BUD20, BUD23, BUD3, BUD32, BUD4, BUD6, BUD7, BZZ1, CAB1, CAB2, CAB4, CAB5, CAC2, CAF120, CAF130, CAF16, CAJ1, CAK1, CAM1, CAN1, CAR1, CAR2, CBC2, CBF1, CBF2, CBF5, CBK1, CBR1, CCA1, CCC1, CCL1, CCR4, CCS1, CCT3, CCT4, CCT5, CCT6, CCW14, CCZ1, CDC10, CDC11, CDC14, CDC16, CDC19, CDC21, CDC24, CDC25, CDC28, CDC3, CDC33, CDC34, CDC39, CDC40, CDC42, CDC48, CDC5, CDC50, CDC53, CDC55, CDC7, CDC9, CET1, CFT1, CFT2, CHO1, CHS2, CHS3, CIK1, CIN1, CIN8, CIS3, CIT1, CIT2, CKA1, CKA2, CKB1, CKB2, CKS1, CLA4, CLB2, CMC2, CMS1, CNB1, COA6, COF1, COG1, COG3, COG6, COG7, COP1, COT1, COY1, CPR1, CPR3, CPR4, CPR5, CPR8, CPT1, CRM1, CRN1, CRP1, CRT10, CRZ1, CSC1, CSG2, CSL4, CSR1, CST26, CST6, CTF18, CTK1, CTK2, CTR1, CTR86, CTS1, CTT1, CUE1, CUE4, CUS1, CUZ1, CWC2, CWC25, CWP1, CYK3, CYR1, CYS3, CYS4, CYT1, DAP1, DAP2, DBF4, DBP2, DBP5, DCP1, DCP2, DCS1, DCS2, DDI1, DDP1, DED1, DEG1, DET1, DFR1, DGA1, DGK1, DGR2, DIA4, DIP5, DIS3, DJP1, DLD3, DNF1, DNF2, DNF3, DNM1, DOA1, DOA4, DOG2, DOP1, DOT5, DOT6, DPB2, DPB4, DPM1, DPP1, DPS1, DRN1, DRS2, DSD1, DSF2, DSK2, DSL1, DUG1, DUG2, DUN1, DUR1,2, DUS3, DUS4, DUT1, DYN1, DYN2, EAF1, EAF3, EAF5, EAP1, EAR1, EBS1, ECM1, ECM15, ECM16, ECM29, ECM3, ECM33, ECM5, ECM7, ECT1, EDC3, EFM1, EFR3, EGD1, EGD2, EHT1, EIS1, ELG1, ELM1, ELP2, ELP3, ELP4, EMC2, EMC3, EMG1, EMI2, EMP24, EMP47, EMP65, EMP70, EMW1, ENA1, ENB1, ENO1, ENO2, ENP1, ENT1, ENT3, ENT5, ENV10, ENV9, EPL1, EPS1, EPT1, ERD1, ERD2, ERG1, ERG10, ERG12, ERG20, ERG27, ERG6, ERG8, ERG9, ERP1, ERP2, ERP3, ERP4, ERP5, ERP6, ERS1, ERV2, ERV25, ESA1, ESBP6, ESC1, ESF2, ESL1, ESL2, ESS1, EXG1, EXG2, EXO70, EXO84, FAA1, FAA2, FAA3, FAA4, FAF1, FAP1, FAP7, FAR11, FAR8, FAS1, FAS2, FAT1, FBA1, FCP1, FCY1, FES1, FET3, FET4, FET5, FIN1, FIP1, FIR1, FIS1, FKH1, FKH2, FKS1, FLC1, FLC2, FLC3, FMP30, FMP42, FMP45, FMP52, FOL1, FOL2, FPK1, FPR1, FPS1, FRD1, FRE6, FRE8, FRK1, FSH1, FSH3, FTH1, FUB1, FUI1, FUM1, FUN12, FUN30, FUS3, FYV8, GAL11, GAL83, GAP1, GAS1, GAS3, GAS5, GAT1, GBP2, GCD7, GCN1, GCN2, GCN20, GCN5, GCS1, GCY1, GDB1, GDH1, GDS1, GDT1, GEA1, GEF1, GET1, GET3, GFA1, GID7, GID8, GIM5, GIN4, GIP3, GIS1, GIS4, GLC7, GLC8, GLE1, GLE2, GLK1, GLN1, GLN4, GLO1, GLO3, GLR1, GMC1, GMH1, GNA1, GND1, GND2, GNP1, GOR1, GOS1, GPA1, GPA2, GPB1, GPB2, GPD1, GPD2, GPI12, GPI15, GPM1, GPP1, GPP2, GPT2, GPX2, GRE2, GRE3, GRH1, GRR1, GRS1, GRS2, GRX1, GRX2, GRX3, GRX4, GRX6, GRX7, GSF2, GSY1, GSY2, GTR1, GTR2, GTT1, GUK1, GUP1, GUS1, GYL1, GYP1, GYP5, HAA1, HAL5, HAL9, HAM1, HAP1, HAT1, HAT2, HBN1, HBT1, HCH1, HCR1, HDA1, HDA2, HDA3, HEH2, HEK2, HEM12, HEM13, HEM2, HEM3, HEM4, HER1, HFA1, HFD1, HFI1, HGH1, HHF1, HIP1, HIR1, HIR2, HIS6, HIT1, HMF1, HMG1, HMO1, HMS2, HNT1, HNT2, HOC1, HOF1, HOG1, HOL1, HOM2, HOM6, HOS3, HOS4, HOT13, HPM1, HPR1, HPT1, HRB1, HRI1, HRK1, HRP1, HRR25, HRT3, HSC82, HSF1, HSH155, HSH49, HSL1, HSL7, HSM3, HSP104, HSP12, HSP150, HSP26, HSP30, HSP31, HSP60, HSP78, HSP82, HST1, HST2, HSV2, HTB1, HTB2, HTS1, HXK1, HXK2, HXT2, HXT7, HYM1, HYP2, HYR1, ICP55, ICS2, IDH1, IDI1, IDP3, IDS2, IES1, IFM1, IGO1, IGO2, IKI3, IKS1, IMD3, IMH1, IML1, IML2, INM1, INM2, INO80, IOC2, IOC3, IPI1, IPP1, IPT1, IRA2, IRC20, IRC25, IRE1, IRR1, ISA1, ISC1, ISN1, ISU1, ITR1, ITR2, IVY1, IWR1, IZH2, IZH4, JAC1, JEM1, JIP4, JJJ1, JJJ2, KAE1, KAP104, KAP114, KAP120, KAP122, KAP123, KAP95, KAR2, KCC4, KCS1, KEI1, KEL2, KHA1, KIN1, KIN2, KIN3, KIN4, KIP1, KKQ8, KOG1, KRE28, KSP1, KTI12, KTR1, KTR3, KTR4, KTR5, KTR6, LAA1, LAM1, LAM6, LAP2, LAP3, LCB2, LCB4, LCB5, LCD1, LCL3, LDB19, LDH1, LEM3, LEU4, LHS1, LIA1, LIP5, LNP1, LOA1, LOC1, LOS1, LPP1, LRE1, LRG1, LRO1, LSB3, LSB6, LSM1, LSM12, LSM2, LSM4, LSM7, LSP1, LST4, LST8, LTP1, LTV1, LYS1, LYS20, LYS21, MAD2, MAD3, MAE1, MAF1, MAK11, MAK3, MAP1, MAP2, MBF1, MCA1, MCD1, MCD4, MCH1, MCH4, MCM2, MCM21, MCM3, MCM4, MCM5, MCM6, MCM7, MDG1, MDH1, MDH2, MDH3, MDJ1, MDJ2, MDM1, MDM35, MDN1, MDS3, MDY2, MEC1, MED1, MED11, MED2, MED4, MED6, MED7, MEF1, MET13, MET18, MET22, MET3, MET30, MET31, MET5, MET6, MET7, MET8, MEU1, MEX67, MFB1, MFT1, MGA2, MGM1, MGS1, MHF1, MHF2, MHR1, MIC19, MIG1, MIG2, MIM1, MIS1, MIT1, MIX23, MKS1, MLC1, MLH1, MLP2, MMF1, MMM1, MMS2, MNN4, MNT3, MOB2, MON2, MOT1, MOT2, MPD1, MPE1, MPP6, MPS3, MRD1, MRE11, MRH1, MRH4, MRI1, MRL1, MRN1, MRPL3, MRS6, MRT4, MRX10, MRX9, MSA1, MSB1, MSB2, MSB3, MSC1, MSC2, MSF1, MSH3, MSH6, MSI1, MSL5, MSN4, MSN5, MSS11, MSS116, MSS4, MST28, MSY1, MTC1, MTC2, MTC4, MTC5, MTD1, MTG1, MTR10, MTR2, MUD1, MUD2, MUM2, MUP1, MVP1, MXR1, MYO1, MYO3, MYO4, MYO5, NAB2, NAB3, NAM7, NAP1, NAR1, NAS6, NAT1, NBP1, NBP35, NCE102, NCL1, NCP1, NCS6, NDC1, NDC80, NEM1, NEO1, NEW1, NFI1, NFS1, NGG1, NGL1, NGL2, NGR1, NHA1, NIF3, NIP7, NIT3, NMA1, NMD3, NMD5, NNF1, NOB1, NOC2, NOC4, NOG1, NOP14, NOP15, NOT3, NOT5, NPL3, NPL4, NPP1, NPR1, NPR2, NPR3, NPT1, NQM1, NRK1, NRM1, NSE3, NSG1, NSR1, NTG1, NTH2, NTO1, NUC1, NUF2, NUM1, NUP133, NUP159, NUP2, NUP60, NUP82, NUS1, NUT1, NVJ1, NYV1, OCH1, OGG1, OPI1, OPI10, OPT1, OPY1, OPY2, ORC2, ORC3, ORC4, ORC5, ORC6, OSH2, OSH3, OSM1, OST5, OXR1, OYE2, PAB1, PAH1, PAI3, PAL1, PAN1, PAN6, PAP2, PAT1, PBI2, PBP1, PBP2, PBP4, PCF11, PCL6, PCM1, PDC1, PDE2, PDH1, PDR1, PDR15, PDR16, PDR5, PDS5, PDX3, PEA2, PEF1, PEP1, PEP4, PEP7, PEX1, PEX10, PEX11, PEX13, PEX14, PEX15, PEX17, PEX19, PEX29, PEX3, PEX30, PEX5, PEX6, PEX8, PFA4, PFF1, PFK1, PFK2, PGA1, PGA2, PGA3, PGC1, PGD1, PGI1, PGK1, PHM7, PHO13, PHO2, PHO4, PHO8, PHO81, PHO84, PHO85, PHO88, PHO91, PHR1, PIB1, PIB2, PIF1, PIH1, PIK1, PIL1, PIN3, PIN4, PIS1, PKC1, PKH3, PLB1, PLB2, PMA1, PMA2, PMC1, PMD1, PMI40, PMR1, PMU1, PNC1, PNG1, PNO1, POA1, POB3, POF1, POL2, POL31, POM152, POM34, POP2, POP4, POP6, POP8, POR1, PPA2, PPH21, PPH3, PPN1, PPT1, PPZ1, PPZ2, PRC1, PRD1, PRE1, PRE10, PRE2, PRE3, PRE4, PRE5, PRE6, PRE7, PRE8, PRE9, PRI1, PRM15, PRM8, PRO2, PRP11, PRP16, PRP18, PRP19, PRP2, PRP21, PRP22, PRP3, PRP38, PRP39, PRP4, PRP42, PRP6, PRP9, PRR1, PRT1, PRX1, PSD2, PSE1, PSF1, PSK1, PSK2, PSP1, PSP2, PSR1, PST1, PST2, PSY4, PTA1, PTC2, PTC3, PTH2, PTI1, PTK1, PTK2, PTM1, PTP1, PTP3, PTR2, PTR3, PUB1, PUN1, PUP1, PUP2, PUP3, PUS4, PUS7, PWP1, PWP2, PYK2, QNS1, QRI1, RAD23, RAD26, RAD27, RAD3, RAD30, RAD4, RAD5, RAD50, RAD51, RAD52, RAD53, RAD59, RAD7, RAI1, RAM1, RAP1, RAS1, RAS2, RAT1, RAX1, RAX2, RBK1, RBS1, RCR1, RCR2, RCY1, RDI1, RDL1, RDS2, RDS3, REF2, REG1, RER1, RER2, RET3, REX3, RFA1, RFA3, RFC3, RFC4, RGD2, RGI1, RGP1, RGR1, RHB1, RHO1, RHO3, RHO4, RHO5, RIB1, RIB3, RIB4, RIM101, RIM15, RIM20, RIM8, RIO1, RIO2, RIX1, RKI1, RKM1, RKR1, RLI1, RMD8, RME1, RMT2, RNA1, RNA14, RNA15, RNH1, RNH201, RNQ1, RNR1, RNR2, RNR4, RNY1, ROD1, ROT2, ROY1, RPA12, RPA135, RPA14, RPA43, RPB11, RPB2, RPB7, RPC10, RPC17, RPC34, RPC37, RPC82, RPL10, RPL14A, RPL16A, RPL16B, RPL21A, RPL22A, RPL24A, RPL24B, RPL25, RPL26A, RPL26B, RPL3, RPL32, RPL33A, RPL36A, RPL36B, RPL37A, RPL37B, RPL38, RPL39, RPL4B, RPL6A, RPL6B, RPL8A, RPL8B, RPL9A, RPL9B, RPM2, RPN1, RPN10, RPN11, RPN14, RPN3, RPN6, RPN8, RPN9, RPO21, RPO26, RPO41, RPP1, RPP2A, RPS12, RPS15, RPS20, RPS29A, RPS29B, RPS31, RPS5, RPT1, RPT2, RPT5, RRD2, RRM3, RRP12, RRP17, RRP43, RSC1, RSC2, RSC30, RSC4, RSF2, RSN1, RSP5, RSR1, RTC1, RTC3, RTC4, RTG3, RTK1, RTN2, RTS1, RTS2, RTT101, RTT103, RUB1, RUP1, SAC1, SAC3, SAC6, SAC7, SAF1, SAK1, SAM3, SAM37, SAM4, SAN1, SAP1, SAP155, SAP185, SAP190, SAP30, SAR1, SAS3, SAS4, SAT4, SAY1, SBA1, SBE22, SBH2, SBP1, SCC2, SCC4, SCH9, SCL1, SCM3, SCP1, SCP160, SCS2, SCS22, SCW10, SCW4, SDC1, SDO1, SDS22, SDS23, SDS24, SDS3, SEC10, SEC12, SEC13, SEC14, SEC16, SEC17, SEC18, SEC21, SEC22, SEC24, SEC26, SEC27, SEC28, SEC3, SEC31, SEC4, SEC5, SEC6, SEC63, SEC65, SEC8, SEC9, SED5, SEG1, SEN1, SER2, SET1, SET2, SET3, SET5, SEY1, SFA1, SFB2, SFH5, SFP1, SFT1, SGF29, SGF73, SGM1, SGT1, SHB17, SHE10, SHM2, SHO1, SHP1, SHQ1, SHS1, SIM1, SIN3, SIN4, SIP1, SIP2, SIP3, SIR2, SIR3, SIR4, SIS1, SIS2, SIT4, SKG1, SKG3, SKG6, SKI8, SKN7, SKO1, SKP1, SKY1, SLA1, SLC1, SLG1, SLI15, SLM1, SLT2, SLU7, SLY1, SLY41, SMC4, SMC6, SMD1, SMD2, SMD3, SMF1, SMM1, SMX3, SNA4, SNF1, SNF12, SNF4, SNF5, SNF6, SNF7, SNG1, SNL1, SNP1, SNQ2, SNT1, SNT309, SNZ1, SOD1, SOG2, SOK2, SOL1, SOL3, SPA2, SPC105, SPC110, SPC25, SPC29, SPC42, SPC72, SPC97, SPE2, SPE3, SPF1, SPO7, SPP381, SPP382, SPP41, SPT14, SPT16, SPT2, SPT20, SPT23, SPT5, SPT6, SQS1, SRB2, SRB4, SRB5, SRB8, SRC1, SRL2, SRM1, SRO7, SRO77, SRO9, SRP1, SRP102, SRP54, SRV2, SRY1, SSA1, SSA2, SSA4, SSB2, SSC1, SSD1, SSE1, SSE2, SSH4, SSL1, SSL2, SSN2, SSO2, SSP120, SSQ1, SSU1, SSZ1, STB1, STB2, STB3, STB6, STE18, STE2, STE20, STE4, STE5, STE6, STM1, STP1, STR2, STR3, STT4, STU1, STU2, SUA5, SUB2, SUC2, SUI1, SUI3, SUR2, SUR7, SVF1, SWC5, SWD1, SWH1, SWI3, SWI4, SWI5, SWI6, SWP1, SWP82, SWR1, SXM1, SYF1, SYF2, SYG1, SYH1, SYO1, SYP1, TAF1, TAF10, TAF11, TAF12, TAF2, TAF3, TAF4, TAF5, TAF6, TAF9, TAH11, TAH18, TAL1, TAN1, TAT1, TAT2, TAX4, TCB1, TCB2, TCO89, TDA1, TDA10, TDA5, TDH1, TDH2, TDH3, TEF1, TEF4, TES1, TEX1, TFA2, TFB1, TFB2, TFB3, TFB4, TFC3, TFC4, TFG1, TFG2, THG1, THP2, THR1, THR4, THS1, TIF1, TIF11, TIF3, TIF34, TIF35, TIF4631, TIF4632, TIF6, TIM22, TIM54, TIP20, TIP41, TKL1, TKL2, TLG2, TMA17, TMA19, TMA64, TMA7, TMN2, TNA1, TOD6, TOF1, TOF2, TOM1, TOM40, TOP2, TOR1, TOR2, TOS1, TOS4, TPA1, TPD3, TPI1, TPK1, TPK2, TPK3, TPM1, TPM2, TPN1, TPO5, TPS2, TPT1, TRE1, TRE2, TRI1, TRM1, TRM10, TRM2, TRM5, TRM82, TRM9, TRP4, TRP5, TRR1, TRS31, TRS33, TRS65, TRS85, TRX1, TRX2, TRZ1, TSC10, TSC11, TSC13, TSR1, TTI1, TUB4, TUM1, TUS1, TVP18, TYS1, TYW1, UBA1, UBA2, UBC1, UBC13, UBC4, UBC6, UBC7, UBP1, UBP10, UBP12, UBP13, UBP14, UBP15, UBP2, UBP3, UBP7, UBP8, UBX2, UBX3, UBX4, UBX5, UFD1, UFD2, UFD4, UFO1, UGA1, UGP1, UIP3, UIP5, ULS1, UME1, UME6, UNG1, UPC2, UPF3, URA2, URA4, URA5, URA6, URA7, URA8, URC2, URH1, UTP10, UTP20, UTP23, UTR1, UTR4, VAC7, VAM3, VAM6, VAM7, VAN1, VAS1, VBA1, VBA4, VHC1, VHS3, VID22, VID27, VID28, VID30, VMA1, VMA10, VMA13, VMA16, VMA4, VMA5, VMA6, VMA7, VMA9, VMS1, VOA1, VPS1, VPS13, VPS15, VPS20, VPS21, VPS33, VPS34, VPS36, VPS4, VPS41, VPS45, VPS51, VPS52, VPS53, VPS54, VPS64, VPS72, VPS74, VPS75, VPS8, VRG4, VRP1, VTA1, VTC1, VTH1, VTI1, VTS1, WAR1, WSC2, WSC4, WTM1, WTM2, WWM1, XDJ1, XRN1, YAE1, YAF9, YAK1, YAP1, YAP1802, YAP3, YAR1, YBT1, YCH1, YCK1, YCK2, YCK3, YCP4, YDJ1, YEF3, YFH7, YHB1, YHC1, YIF1, YIM1, YIP3, YJU3, YKE2, YKT6, YLF2, YNG2, YNK1, YPD1, YPK1, YPK3, YPK9, YPP1, YPQ1, YPQ2, YPR1, YPT1, YPT10, YPT11, YPT31, YPT32, YPT52, YPT6, YPT7, YRA1, YRB1, YRB2, YRB30, YRO2, YSA1, YSH1, YSP2, YTA12, YTA6, YTA7, YTH1, YUR1, YVH1, ZEO1, ZPR1, ZPS1, ZRC1, ZRT1, ZRT2, ZTA1, ZUO1
